# The ISW1 and CHD1 chromatin remodelers suppress global nucleosome dynamics in living yeast cells

**DOI:** 10.1101/2025.04.17.649351

**Authors:** Zhuwei Xu, Hemant K. Prajapati, Peter R. Eriksson, David J. Clark

## Abstract

The budding yeast genome is globally accessible to DNA methyltransferases in living cells, unlike in isolated nuclei, where it is mostly inaccessible. Here, we assess the roles of the RSC, ISW1 and CHD1 ATP-dependent chromatin remodelers in generating nucleosome dynamics in vivo. We compare DNA methylation rates in wild type cells and chromatin remodeler mutants by normalizing nuclear methylation rates to the non-nucleosomal mitochondrial DNA methylation rate in each strain. Depletion of subunits of the RSC, ISW1 or CHD1 ATP-dependent chromatin remodelers has little effect on normalized methylation rate. A decaying sine wave model used to fit nucleosome phasing data shows that nucleosome dynamics decrease with distance from the promoter in an Isw1/Chd1-dependent manner. Depletion of both Isw1 and Chd1 increases the methylation rate, suggesting that these remodelers act together to suppress nucleosome dynamics. Furthermore, the TFIIIB and TFIIIC transcription factors exhibit differential dynamics at tRNA genes. Our analysis provides insight into nucleosome and transcription factor dynamics in vivo.

**Teaser:** The budding yeast ISW1 and CHD1 ATP-dependent remodelers organize chromatin on genes, slowing nucleosome dynamics in living cells.

## Introduction

The structural subunit of chromatin is the nucleosome core, which contains ∼147 bp of DNA wrapped in 1.7 negative superhelical turns around a central core histone octamer composed of two molecules each of the four core histones (H2A, H2B, H3 and H4) (*1*). In vitro, the nucleosome is very stable; extreme salt concentrations are required to dissociate the histones from the DNA (*2*). In less extreme conditions, but still non-physiological, nucleosomes can slide along DNA without dissociation (*3, 4*). Although the nucleosome is a stable entity, the terminal 10-20 bp are much less strongly bound, undergoing rapid cycles of dissociation and re-binding on a seconds time scale, providing access to the terminal DNA (*5, 6*). These observations are based on in vitro experiments using native chromatin fragments or reconstituted chromatin. This rather static view of chromatin accounts for its generally repressive nature in vitro. Current models of gene regulation are based on this intrinsic repressive property of chromatin.

Many studies have focused on the question of how sequence-specific transcription factors access their binding sites in chromatin. The thermodynamic stability of the nucleosome contrasts with the transient and reversible nature of transcription factor binding in vitro and in vivo. Thus, a simple binding competition model is untenable, unless there are several transcription factor binding sites within the same nucleosome. In this case, histone displacement can occur if the transcription factor concentrations are high enough (*7*). The pioneer transcription factors are an alternative solution to the problem (*8*). They bind their cognate sites within nucleosomes with high affinity, without displacing the nucleosome (*9*). These considerations, together with the fact that transcription is severely inhibited by chromatin in vitro, led to the discovery of the ATP-dependent chromatin remodelers, such as NURF and SWI/SNF (*10, 11*). In budding yeast, SWI/SNF and the similar RSC complex use the free energy of ATP hydrolysis to move, remove, or conformationally alter nucleosomes (*12–15*), whereas ISW1 and CHD1 are nucleosome spacing enzymes (*16–18*). In vitro, remodelers can convert a static chromatin structure into a dynamic structure. Indeed, this is the basis of a standard assay for remodeling activity, in which chromatin is digested with a restriction enzyme in the presence or absence of the remodeler and ATP.

The accessibility of chromatin in isolated yeast nuclei is severely limited (*19–21*). Previously, we incubated nuclei with the restriction enzyme AluI (*19*) or with the *E. coli dam* sequence-specific DNA methyltransferase (Dam) (*20*). We found that 65-75% of yeast genic DNA is inaccessible. This result is expected if nucleosomes block access to the enzyme probe. However, using inducible expression of Dam in yeast cells, we showed that virtually all of the yeast genome is accessible in vivo, unlike in isolated nuclei (*20*). Only the point centromeres and, to a lesser extent, the silenced mating type loci (*HMLa* and *HMRa*), are inaccessible in living yeast cells. We proposed that chromatin in isolated yeast nuclei is essentially static, whereas chromatin in living cells is highly dynamic (*20*). We also showed that the RSC, ISW1 and CHD1 ATP-dependent chromatin remodelers contribute to these dynamics in vivo. The highly dynamic nature of yeast chromatin in vivo suggests that nucleosomes are not necessarily blocking access to the genome, but might instead be packaging it without maintaining it in a repressed state. More recently, we have shown that nucleosomes are also globally dynamic in human cells, in both euchromatin and heterochromatin (*22*).

Our current aim is to begin to determine the roles of the various yeast chromatin remodelers in generating nucleosome dynamics in vivo. Here, we present an in depth analysis of the contributions of the RSC, ISW1 and CHD1 chromatin remodelers to nucleosome dynamics in vivo, based on our published yeast data (*20*). We compare methylation rates in wild type (WT) cells and remodeler mutants in vivo using mitochondrial DNA (mtDNA) as a normalization control. We find that the genic methylation rate increases after depletion of both the ISW1 and CHD1 remodelers, indicating that together, they suppress nucleosome dynamics. We use a decaying sine wave model to fit nucleosome phasing data and to obtain phasing data for single genes in vivo. We show that nucleosome dynamics decrease with distance from the promoter and that this effect is also dependent on ISW1 and CHD1. Finally, we dissect the dynamics of the TFIIIB and TFIIIC transcription factors at tRNA genes in vivo.

## Results

In our previous experiments, we incubated purified yeast nuclei with increasing amounts of Dam to methylate accessible GATC sites and then purified genomic DNA for qDA-seq (*20*). Briefly, this technique involves digestion of genomic DNA with the restriction enzyme DpnI, which is specific for GATC sites methylated on both strands (m^6^A). DpnI cuts to give blunt ends (ending with GA and beginning with TC), although some molecules lose the terminal ‘A’ or ‘T’. The DpnI-digested DNA is then fragmented by sonication and subjected to paired-end sequencing. To quantify the fraction methylated for each of the ∼36,000 GATC sites in the yeast genome, we counted the number of DNA molecules for each site that end on the G or the A in GATC and divided by the fragment coverage of the G. Similarly, we counted the number of DNA molecules that begin with the T or the C and divided by the fragment coverage of the C. This ratio is the fraction methylated by Dam.

We measured DNA accessibility in vivo by expression of Dam with a C-terminal auxin-dependent degron and three HA tags, using an inducible promoter driven by a single high-affinity binding site for the Gcn4 transcription activator (*20*). Auxin was added before induction to reduce background Dam expression. Activation by Gcn4 was induced by addition of sulfometuron methyl (SM) (*23*). Depletion of the Rsc8 structural subunit of RSC, or of the Isw1 ATPase subunit of ISW1, or of the Chd1 ATPase, was achieved by attaching the same auxin-dependent degron and three HA tags to the endogenous gene. Cells were treated with auxin for 4 h to deplete the remodeler and to reduce background Dam expression, and then SM was added to induce Dam for 4 h in the continued presence of auxin. Genomic DNA was purified at various times and subjected to qDA-seq. In other experiments, we used the M.SssI DNA methyltransferase to methylate C in CG in vivo, purified the genomic DNA and subjected it to nanopore long-read sequencing to detect m^5^CG. The M.SssI data have much higher resolution than the Dam data because there are many more CG sites than GATC sites in the yeast genome (∼355,000).

### Normalization of Dam methylation rates in remodeler mutants using mtDNA

Previously, we did not compare methylation rates in remodeler mutants with wild type cells because different strains produced different amounts of Dam and with different kinetics (*20*). We showed that, in wild type cells, the amount of Dam protein peaks at about 150 min after induction and then decreases, whereas Dam protein slowly increases in cells being depleted of Rsc8, producing about half as much Dam as in wild type cells.

Isw1-depleted cells produce similar amounts of Dam with similar kinetics to wild type cells, whereas more Dam is produced in Chd1-depleted cells than in wild type cells (Fig. S1). Cells depleted of both Isw1 and Chd1 produce two to three times more Dam than wild type cells, with similar kinetics (*20*). This variation is a critical issue, because the methylation rate is expected to be directly proportional to the enzyme concentration at time ‘*t*’.

An internal control is needed to normalize for Dam concentration in the various mutants. Unexpectedly, we discovered that mtDNA is also methylated by Dam in vivo (Fig. 1A). It appears that Dam (a prokaryotic protein) is transported into yeast mitochondria. In wild type cells, GATC sites in mtDNA are methylated more slowly than those in promoter NDRs but slightly faster than GATC sites in gene bodies (Fig. 1B). We obtained apparent rate constants for mtDNA, gene bodies and promoter NDRs by assuming pseudo-first order rates and plotting the log of the unmethylated fraction against time of SM treatment (Fig. 1C). The assumption is validated by the observed straight lines with *R^2^*values ranging from 0.86 to 0.99 for all biological replicates (Table S1). Since mtDNA is not assembled into nucleosomes, it is unlikely to be directly affected by chromatin remodelers. Instead, the mtDNA methylation rate might be determined by the rate at which Dam enters the mitochondria, and/or by the distribution of Abf2, a mitochondrial DNA-binding protein (*24*).

**Fig. 1.**
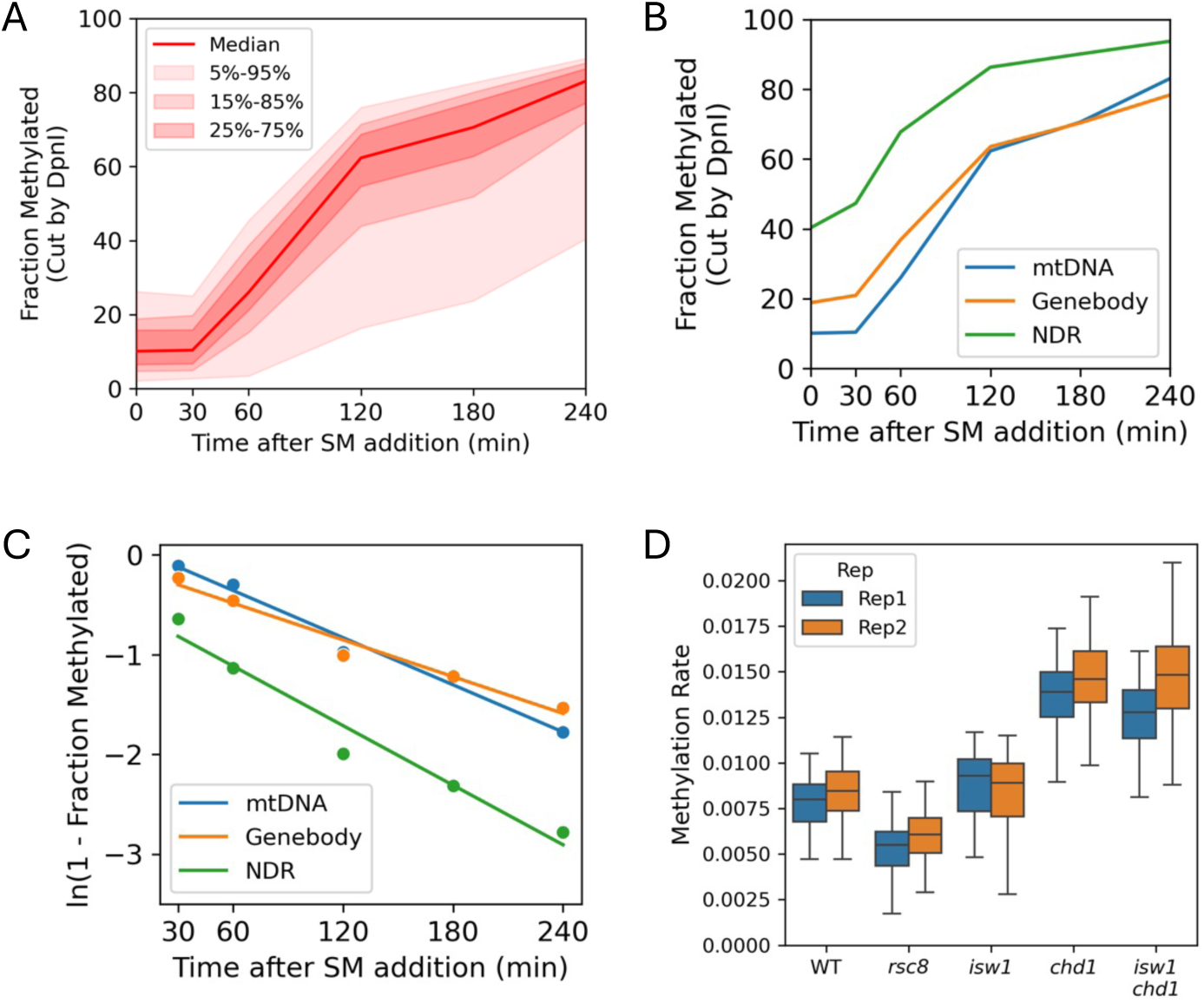
Dam methylation of mtDNA in living cells. (**A**) Time course of Dam methylation in mitochondria after SM induction. Red line and shading: median GATC methylation with data range shown. (**B**) Time course of median Dam methylation of mtDNA, gene bodies and promoter NDRs after SM induction. (**C**) Dam methylation rate of the median GATC site in mtDNA, gene bodies and promoter NDRs. (**D**) Methylation rates of mtDNA in cells depleted of remodeler subunits (data are shown for biological replicates). The distribution of methylation rates for individual GATC sites in mtDNA are illustrated in box plots (the box contains 25%-75% of the data; the line is the median; the whiskers represent 1.5 times the inter-quartile range to the farthest data points).

We used the GATC sites in the circular 85.8 kb yeast mtDNA as an internal control for methylation rate comparisons between strains. We calculated the apparent rate constant for each GATC site in mtDNA in wild type cells, Rsc8-depleted cells, Isw1-depleted cells, Chd1-depleted cells and cells depleted of both Isw1 and Chd1. As expected from the variation in Dam concentration in the remodeler mutants, we observed a fairly wide range in rate constant distributions (Fig. 1D). The biological replicate experiments are consistent with one another. Rsc8-depleted cells methylate their mtDNA more slowly than wild type cells, whereas cells depleted of Chd1 or both Chd1 and Isw1 methylate their mtDNA much faster. Isw1-depleted cells methylate their mtDNA slightly faster than wild type. These observations are qualitatively consistent with the Dam protein levels observed in these strains (*20*) (Fig. S1). We attribute these differences in mtDNA methylation rate to the variation in Dam concentration.

### Isw1 and Chd1 together suppress nucleosome dynamics on genes in vivo

We computed the apparent rate constant for every GATC site in the nuclear genome and divided it by the median methylation rate constant for GATC sites in mtDNA obtained from Fig. 1C (Table S1). We used these ratios to construct a nucleosome phasing plot for all yeast genes in wild type cells (Fig. 2A). The x-axis represents the distance of the GATC site from the dyad of the first (+1) nucleosome on each gene (determined using MNase-seq data (*18*)). The y-axis is the ratio of the apparent rate constant for a GATC site to the median mtDNA site, for all GATC sites in 5398 yeast genes aligned on the dyad of the first (+1) nucleosome. The mean relative methylation rate for gene bodies is defined for phasing plots as the region from -50 to +1000 bp relative to the +1 nucleosome dyad. The mean rate for wild type cells is somewhat slower than that for mtDNA (∼80% of the mtDNA rate, which is set at 1; Fig. 2). The replicates are almost identical after normalization to mtDNA, supporting the use of mtDNA as an internal control. The peaks in the plot represent the average methylation rate for GATC sites that are in the linker DNA between nucleosomes in most cells; the troughs represent the average methylation rate for GATC sites that are inside nucleosomes in most cells (Fig. 2A). GATC sites at the promoter peak, which are generally nucleosome-depleted, are methylated ∼1.3 times faster than GATC sites in mtDNA and ∼1.6 times faster than the average GATC site in gene bodies.

**Fig. 2.**
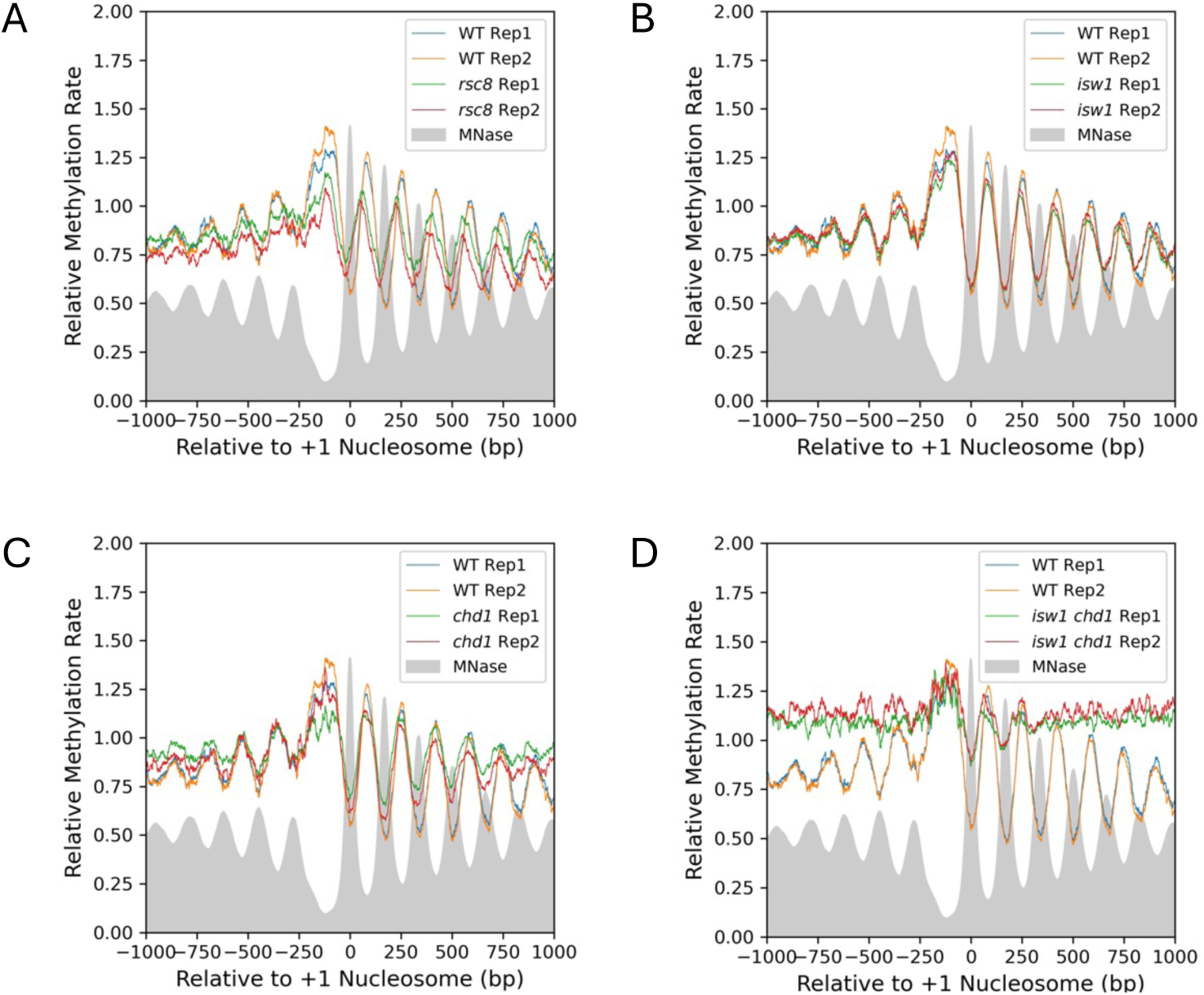
Comparison of normalized average Dam methylation rates in vivo across genes: wild type vs. cells depleted of remodeler subunits. The methylation rate was calculated for each GATC site in the yeast genome and normalized to the internal median mtDNA methylation rate (set to 1). The normalized rates are plotted relative to the location of the +1 nucleosome dyad of 5398 genes (smoothed with a 21-bp window). Grey profile: MNase-seq nucleosome dyad data for wild type cells (*18*) for comparing phasing patterns (arbitrarily normalized to 0.5). (**A**) WT vs. Rsc8-depleted cells. (**B**) WT vs. Isw1-depleted cells. (**C**) WT vs. Chd1-depleted cells. (**D**) WT vs. cells depleted of both Isw1 and Chd1.

In Rsc8-depleted cells, GATC sites predominantly located in linkers are methylated more slowly than linkers in wild type cells, but GATC sites predominantly in nucleosomes are methylated faster than in wild type cells (Fig. 2A). This can be explained by weaker phasing in Rsc8-depleted cells, such that there is a broader distribution of alternative nucleosome positions around the average position. GATC sites in linkers are more likely to be nucleosomal after Rsc8 depletion, and GATC sites in nucleosomal regions are more likely to be in a linker after Rsc8 depletion. We observe a shift of the nucleosomal array on each side of the NDR towards the promoter after Rsc8-depletion (expected from MNase-seq data (*25, 26*)). However, the average methylation rates across genes in Rsc8-depleted and wild type cells are similar. In conclusion, loss of RSC does not affect the overall methylation rate in vivo, with the caveat that Rsc8 is not fully depleted until the end of the time course (*20*).

Depletion of either Isw1 or Chd1 has little effect on average methylation rate (Fig. 2B,C). Loss of Isw1 results in shorter nucleosome spacing with no change in the average +1 nucleosome position, with weaker phasing, as observed in nuclei using MNase-seq (*18*). Similarly, loss of Chd1 results in weaker phasing with little change in average nucleosome positions, again as observed in nuclei using MNase-seq (*18*). However, when both Isw1 and Chd1 are depleted, the methylation rate increases markedly across the first 1 kb of genes (∼1.4 times faster than wild type; Fig. 2D). The nucleosomal arrays are almost completely disrupted; only the +1 nucleosome remains relatively well positioned (*20*). Such disorder was first observed in nuclei from the *isw1Δ chd1Δ* double mutant using MNase-seq (*17*). The promoter NDRs are methylated at similar rates in wild type and doubly depleted cells; the increased rate is observed only over gene bodies, where nucleosomes are located (Fig. 2D). We conclude that Isw1 and Chd1 act together to reduce nucleosome dynamics on genes.

We compared relative methylation rates for GATC sites in different genomic regions in the various mutants. Relative rates for GATC sites in promoter NDRs are similar in wild type and the four mutants (Fig. S2A). However, relative rates for sites in gene bodies are faster in the Isw1/Chd1 depletion double mutant than in wild type and the other mutants (Fig. S2B). Methylation rates at tRNA genes are similar to those at promoter NDRs; they are unaffected by the remodeler mutations, possibly because both are nucleosome-depleted (Fig. S2C). Replication origins (*ARS* elements) are methylated faster than gene bodies but not as fast as tRNA genes and promoters, possibly reflecting the presence of an NDR within each element, corresponding to the binding of the origin recognition complex (Fig. S2D). Ty transposable elements and telomeric regions are methylated at similar rates to gene bodies, but only Ty elements are methylated faster in the Isw1/Chd1 depletion double mutant (Fig. S2E,F). In conclusion, gene bodies and Ty elements are methylated faster in cells depleted of both Isw1 and Chd1 than in wild type cells or the single mutants, probably because they are completely assembled into nucleosomes and therefore more affected by loss of these remodelers.

### Decaying sine wave model for quantifying nucleosome phasing in vivo

We found that a decaying sine wave model provides a quantitative fit to the nucleosome phasing methylation rate profiles (Fig. 3; Table 1). Rate constants for individual GATC sites are plotted (blue dots) relative to the +1 nucleosome dyad; the sine wave fit is shown as a red line. The best fit was obtained by assuming that the sine wave decays exponentially and by adding a linear regression term to model the slope in the data. The degree of fit is described by the coefficient of determination (*R^2^*) adjusted for the number of parameters used for the fit. The wavelength corresponds to the average nucleosome spacing. The initial amplitude corresponds to the trough representing the +1 nucleosome dyad. The amplitude decays exponentially down the gene, described by the ‘Decay’ over one wavelength. The ‘Slope’ represents the change in mean methylation rate over 1 kb. The ‘Adj. Mean Rate’ refers to the mean fitted methylation rate constant normalized to mtDNA in the range -50 to +1000 bp relative to the +1 nucleosome dyad.

**Fig. 3.**
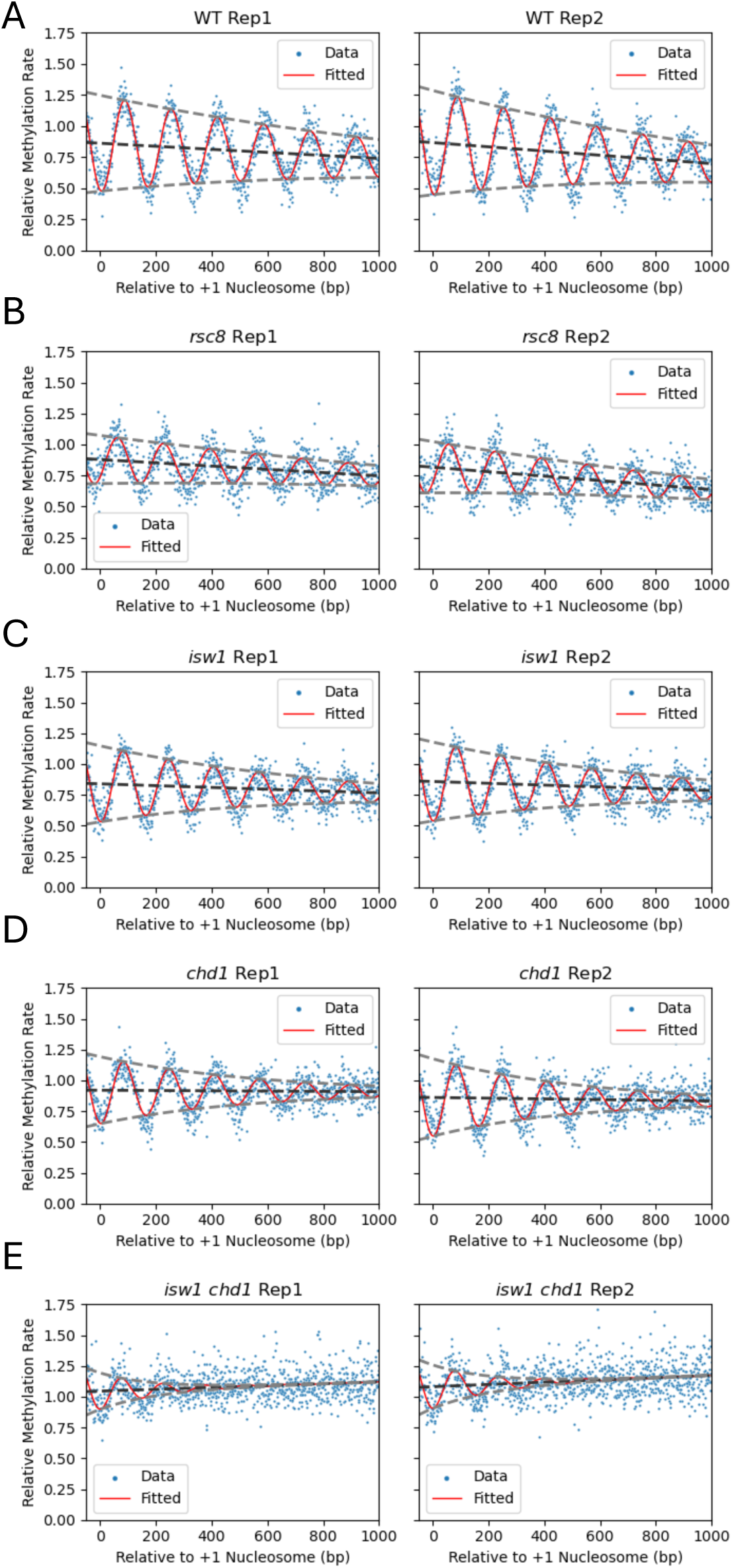
Decaying sine wave model fit to Dam methylation data for wild type and remodeler-depleted cells in vivo. Sine wave analysis provides a quantitative comparison for wild type cells (**A**) and remodeler mutants (**B-E**) (see Table 1 for fit parameters). The methylation rate was calculated for each GATC site in the yeast genome and normalized to the internal median mtDNA methylation rate (set at 1). The normalized rates are plotted relative to the location of the +1 nucleosome dyad of 5398 genes. Values for individual GATC sites are indicated by blue dots. The decaying sine wave fit is shown as a red line. The baseline of the sine wave is indicated by the dark grey dashed line; the decaying amplitudes are indicated by light grey dashed lines.

**Table 1.** Decaying sine wave model parameters fitted to Dam data for wild type and remodeler degron mutants in vivo. The degree of fit is described by the coefficient of determination (*R^2^*) adjusted for the number of parameters used for the fit. The average nucleosome spacing corresponds to the wavelength. The amplitude corresponds to the trough representing the +1 nucleosome dyad. The amplitude decays exponentially down the gene, described by the ‘Decay’ over one wavelength. The ‘Slope’ represents the change in mean methylation rate over 1 kb. The ‘Adj. Mean Rate’ refers to the fitted mean methylation rate constant relative to mtDNA for the first kilobase of all genes.

| Strain | Replicate | Spacing (bp) | Amplitude | Slope per kb | Decay per Period | Adjusted Mean Rate | Adjusted $R^2$ |
| --- | --- | --- | --- | --- | --- | --- | --- |
| WT | 1 | 166.3 | 0.39 | -0.12 | 0.86 | 0.80 | 0.79 |
| WT | 2 | 165.9 | 0.42 | -0.17 | 0.84 | 0.78 | 0.79 |
| *rsc8* | 1 | 165.6 | 0.19 | -0.13 | 0.87 | 0.81 | 0.53 |
| *rsc8* | 2 | 165.8 | 0.21 | -0.18 | 0.86 | 0.72 | 0.59 |
| *isw1* | 1 | 161.6 | 0.31 | -0.07 | 0.80 | 0.80 | 0.71 |
| *isw1* | 2 | 162.0 | 0.32 | -0.07 | 0.80 | 0.81 | 0.71 |
| *chd1* | 1 | 162.7 | 0.27 | -0.01 | 0.75 | 0.91 | 0.57 |
| *chd1* | 2 | 163.5 | 0.31 | -0.03 | 0.73 | 0.84 | 0.58 |
| *isw1 chd1* | 1 | 149.4 | 0.15 | 0.08 | 0.46 | 1.08 | 0.15 |
| *isw1 chd1* | 2 | 156.1 | 0.18 | 0.09 | 0.48 | 1.12 | 0.17 |

### The methylation rate decreases with distance from the promoter in vivo

All model parameters are in good agreement for the wild type biological replicate experiments (Fig. 3A; Table 1). The spacing is 166 bp, which is the same as in nuclei (measured using MNase-seq data (*18*)). The decay in amplitude with distance from the promoter is a measure of the gradual weakening of phasing, which can be explained by increasing adoption of alternative positions relative to the major position, resulting in broader peaks and troughs. This reflects variation in linker length according to the ‘10*n* +5 bp’ rule (*27, 28*), as well as the combined effects of the various chromatin remodelers (*18, 29*).

The downward slope in the sine wave fit is surprising (Fig. 3A). It indicates that the average methylation rate decreases with distance from the promoter; the values represent the change in rate per kb (a slope of 0 would indicate that the peaks and troughs are decaying at the same rate). Although linker methylation (represented by the peaks) decreases markedly along genes, nucleosomal methylation (represented by the troughs), increases only moderately along genes, resulting in a net negative slope. The downward slope is not due to the fact that we are plotting apparent rate constants, because it is also observed in more standard phasing plots of fraction methylated against distance from the +1 nucleosome dyad at different time points (Fig. S3A). A potential explanation is that linker length decreases with distance from the +1 nucleosome. However, this is not the case because the sine wave fit is good and requires a fixed wavelength and therefore a fixed mean linker length (fixed nucleosome spacing). Modelling of each time point for wild type cells shows that the downward slope ranges from ∼3% to ∼6% methylation per kb in vivo, depending on the time point (Fig. S3A). Nuclei show a weaker downward slope, corresponding to a decrease of ∼2% to 3% per kb (Fig. S3B). The slope suggests that promoter-proximal nucleosomes differ from promoter-distal nucleosomes in their dynamics or histone modifications (discussed below).

### Both Chd1 and Isw1 contribute to decreasing methylation rate along genes in vivo

Rsc8-depleted cells differ substantially from wild type only in their phasing amplitude, which is reduced by about half (Fig. 3B; Table 1). The much weaker phasing indicated by the lower amplitude may be explained by genic variation in the degree of shift of the nucleosomal array towards the promoter (*26*). In Isw1-depleted cells, the amplitude is less than in wild type cells and the spacing is shorter (162 bp) (Fig. 3C; Table 1), though not as short as measured by MNase-seq in nuclei from *isw1D* cells (159 bp) (*18*). Chd1-depleted cells also display weaker phasing than wild type cells, as well as slightly shorter spacing (∼163 bp; Fig. 3D; Table 1), consistent with MNase-seq data for *chd1D* cells (164 bp) (*18*). The slope of the sine wave decay is reduced in Isw1-depleted cells and almost absent in Chd1-depleted cells (Table 1), indicating that both Isw1 and Chd1 contribute to the decrease in methylation rate with distance from the promoter.

In cells depleted of both Isw1 and Chd1, the data fit is poor (Table 1) due to the major disruption in genic chromatin organization (Fig. 3E). The spacing is much shorter than wild type, although this value is likely to be inaccurate given the weak phasing and the poor fit (Table 1). We note that we did not measure spacing using MNase-seq previously, because the phasing was too irregular (*18*). The mean methylation rate is faster, at 1.1 times that of mtDNA (wild type is 0.8 times that of mtDNA). The decay slope is positive rather than negative, indicating that the methylation rate *increases* with distance from the promoter in the absence of both Isw1 and Chd1. In conclusion, Isw1 and Chd1 are required for the decreasing methylation rate along genes.

### Genes with longer nucleosome spacing are methylated faster in vivo

Previously, we also measured yeast genome accessibility in vivo by inducible expression of the M.SssI DNA methyltransferase (*20*). This enzyme converts CG to m^5^CG, which we detected using long-read nanopore sequencing. Since CG sites occur at much greater frequency in the yeast genome than GATC sites, M.SssI data have much higher resolution than Dam data. We applied our sine wave model to the apparent rate constant data derived from the M.SssI induction time course (Fig. 4A). The adjusted *R^2^* values are > 0.9 for both replicates, indicating a much better fit than for the Dam data for wild type cells (compare Figs. 3A and 4A; Tables 1 and 2), presumably reflecting the increased resolution. The downward slope in the phasing decay observed in the Dam data is also apparent in the M.SssI data (cf. Fig. 3A) and in phasing plots of the methylated fraction after induction for 4 h (Fig. S4).

**Fig. 4.**
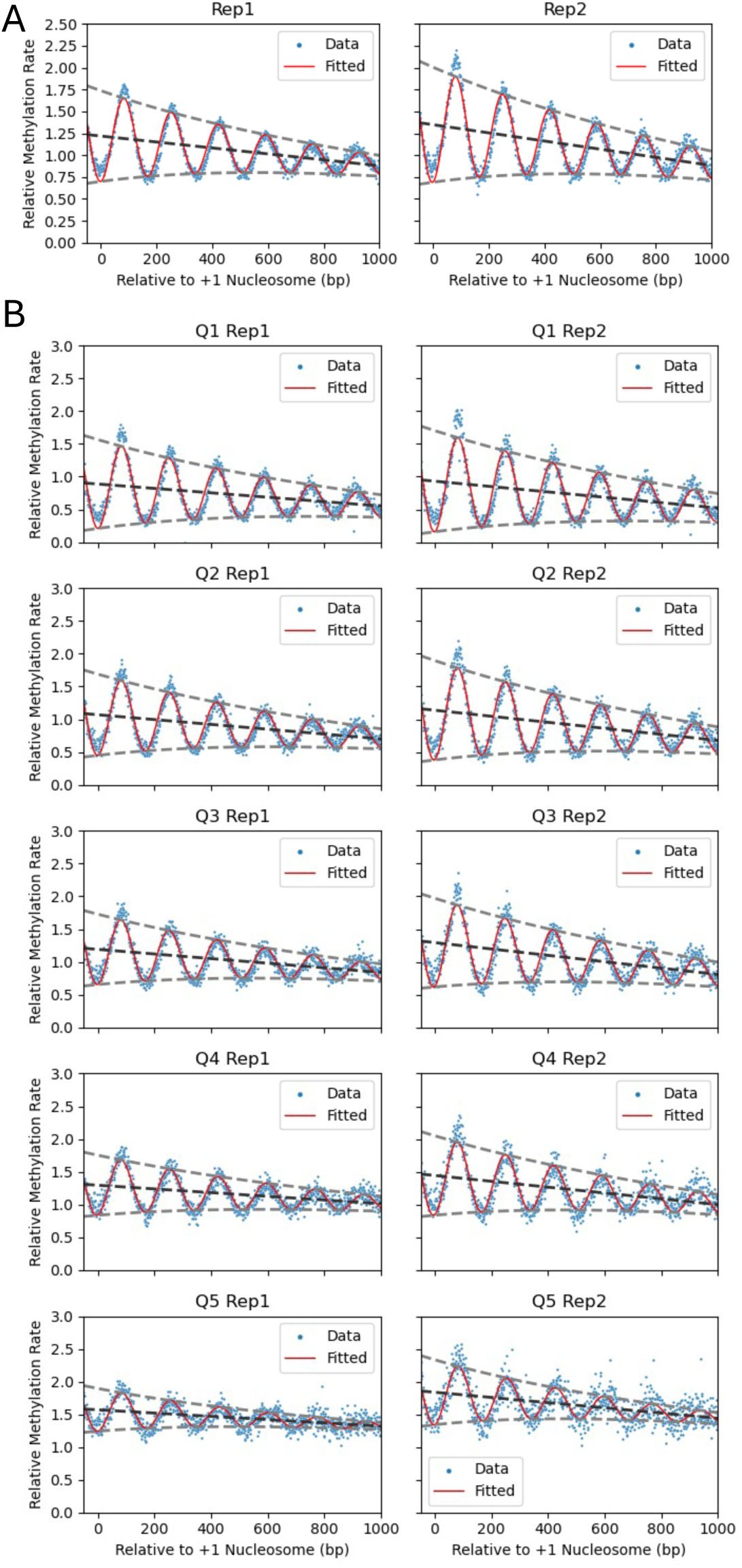
Decaying sine wave model fit to M.SssI methylation nanopore data for wild type cells in vivo. (**A**) Analysis of all genes. The methylation rate was calculated for each CG site in the yeast genome normalized to the genome median rate (set to 1). Individual CG sites are indicated by blue dots. The decaying sine wave fit is shown as a red line. The baseline of the sine wave is indicated by the dark grey dashed line; the decaying amplitudes are indicated by light grey dashed lines. (**B**) Analysis of genes sorted into quintiles using their calculated individual unbiased average methylation rates. Quintile 1 (Q1) contains the slowest methylating genes. See Table 2 for fit parameters.

**Table 2.**
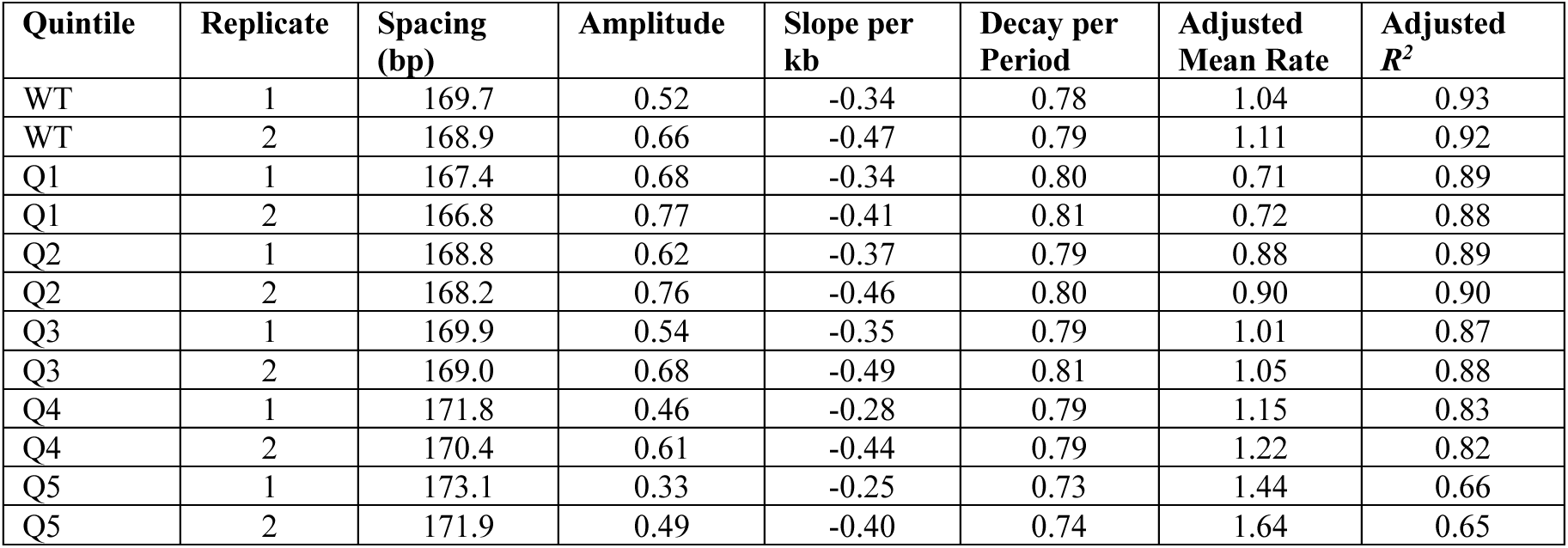
Decaying sine wave model parameters fitted to M.SssI nanopore data for wild type cells in vivo. For all genes and for individual genes grouped by methylation rate (quintiles 1-5; Q5 contains the fastest methylating genes). Parameters are as described in the legend to Table 1.

The nucleosome spacing estimated from the M.SssI data (169-170 bp) is a little higher than expected from MNase-seq data for isolated nuclei (166 bp (*18*)) and from Dam data for living cells (166 bp; Fig. 3A). This difference may result from the fact that the M.SssI data are long read data, whereas MNase and qDA-seq data are short read data. Long read data include a correlation between the positions of neighboring nucleosomes on the same DNA molecule, whereas MNase-seq data give the average for single nucleosomes separated from their neighbors by MNase. Thus, the M.SssI long read data may give the most accurate measure of nucleosome spacing.

The much larger number of methylation target sites for M.SssI allowed us to estimate the mean methylation rate for individual yeast genes in vivo. The simple mean of methylation rates for CG sites in each gene is biased because they are not evenly distributed along the gene. To correct for this bias, we calculated the methylation rate at each CG site from -50 to +1000 relative to the +1 nucleosome dyad for each of 5398 genes and fitted the decaying sine wave model to the data for each gene. The mean methylation rate for each gene was estimated as the mean deviation of the CG sites in each gene from the fitted curve for all genes. For example, if the methylation rates for the CG sites in a particular gene are all above the fitted curve for the population average, this gene is methylated faster than the average.

The 5398 genes were divided into quintiles (Q1 to Q5) according to their interpolated mean methylation rate constant (see Methods). The sine wave model was applied again to the combined data for the genes in each quintile (Fig. 4B; Table 2). The mean rate constant for each quintile increased as expected from the sort (Table 2). Thus, the genes in quintile 5 are methylated ∼2.2 times faster than those in quintile 1. All quintiles show similar decay slopes. The genes in quintile 5 have the weakest phasing (as shown by the amplitudes; Table 2) and the longest spacing (Table 2). The average nucleosome spacing on genes in each quintile increases with methylation rate (ranging from 167 to 172 bp; Table 2). This observation provides an explanation for the differences in average methylation rates: increased spacing indicates longer linkers, which are methylated faster than nucleosomal DNA, resulting in higher average methylation rates. We note that even a change of 5 bp in nucleosome spacing may be important, because it affects the folding trajectory of a chromatin fibre (e.g., (*30*)).

We also observed that the genes in each methylation rate quintile differ substantially in average length, such that longer genes tend to be methylated more slowly than shorter genes (Fig. S5). This observation is consistent with the observation that the promoter-proximal regions of genes are methylated faster than promoter-distal regions (Fig. 4B).

### Methylation rate is not strongly correlated with transcription level

Heavily transcribed genes are associated with weak phasing, irregular spacing and reduced nucleosome occupancy in yeast (*31–35*). We determined whether methylation rate is correlated with transcription level, using RNA-seq data for SM-induced cells (*36*). The transcript levels of the genes in each methylation rate quintile are presented as box plots (Fig. S6). Although the median gene expression level increases slightly and reproducibly with methylation rate, the difference is subtle. The fastest methylating genes are somewhat enriched in highly transcribed genes, since the box is skewed to higher transcript levels, but transcription level is only a minor contributor to methylation rate.

### The fastest methylating genes lack phasing in Isw1/Chd1-depleted cells

To determine the contributions of the ISW1 and CHD1 nucleosome spacing enzymes to the observed genic variation in nucleosome phasing (Fig. 4B), we plotted the Dam methylation rate data for the quintiles of genes identified using high-resolution M.SssI data (Fig. 5; biological replicate data are shown in Fig. S7). The Dam data for wild type cells show the same trends that we observed in the M.SssI data for wild type cells: the mean genic methylation rate increases from Q1 to Q5 (Table 3), the phasing amplitude decreases from Q1 to Q5, and all quintiles display a negative slope (Fig. 5A; Table 3). The sine wave fit is not as good as for the M.SssI data, presumably because there are many fewer GATC sites than CG sites.

**Fig. 5.**
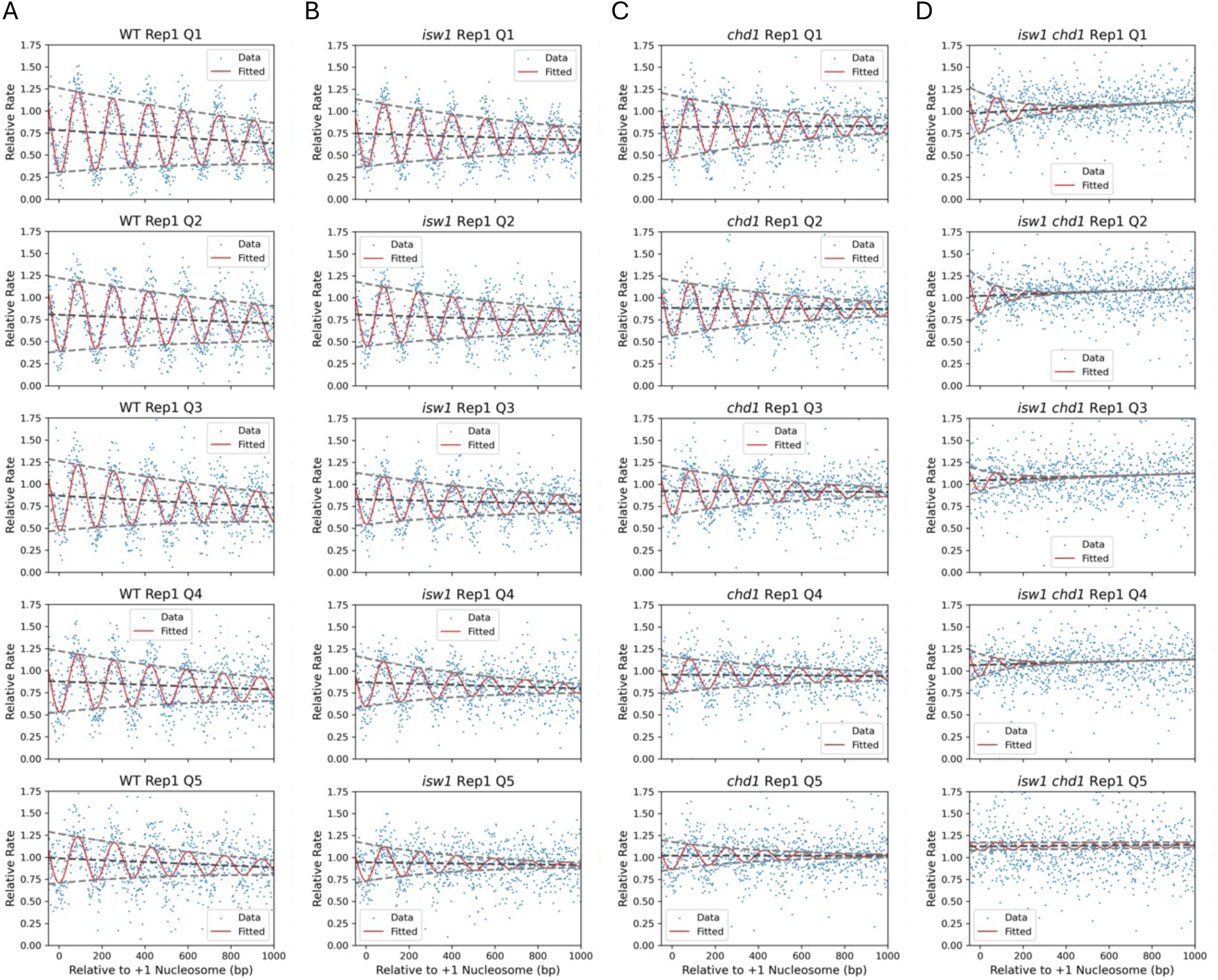
Decaying sine wave model fit to Dam methylation data for wild type and remodeler-depleted cells in vivo: Comparison of methylation rate quintiles. (**A**) wild type; (**B**) Isw1-degron mutant; (**C**) Chd1-degron mutant; (**D**) Isw1-degron and Chd1-degron mutant. See Table 3 for fit parameters. Dam methylation rate data normalized to mtDNA (set to 1) are plotted for each quintile of genes defined using M.SssI nanopore data for replicate 1 (Fig. 4B). Values for individual GATC sites are indicated by blue dots. The methylation rates for each set of genes are plotted relative to the location of the +1 nucleosome dyad. The decaying sine wave fit is shown as a red line. The baseline of the sine wave is indicated by the dark grey dashed line; the decaying amplitudes are indicated by light grey dashed lines. Data for biological replicate 2 are shown in Fig. S7.

**Table 3.** Decaying sine wave model parameters fitted to Dam data for wild type and depletion mutants in vivo. For the quintile data in Fig. 5 (Replicate 1): individual genes grouped by methylation rate (Q5 contains the fastest methylating genes). Parameters are as described in the legend to Table 1. Spacing values are not provided for the Isw1/Chd1 double depletion mutant because the fit is too poor. See Table S2 for the fit to biological replicate 2.

| Strain | Quintile | Spacing (bp) | Amplitude | Slope per kb | Decay per Period | Adjusted Mean Rate | Adjusted $R^2$ |
| --- | --- | --- | --- | --- | --- | --- | --- |
| WT | 1 | 164.2 | 0.48 | -0.15 | 0.89 | 0.70 | 0.65 |
| WT | 2 | 164.1 | 0.42 | -0.10 | 0.89 | 0.75 | 0.56 |
| WT | 3 | 166.7 | 0.39 | -0.13 | 0.86 | 0.80 | 0.44 |
| WT | 4 | 169.9 | 0.34 | -0.09 | 0.85 | 0.83 | 0.35 |
| WT | 5 | 171.5 | 0.28 | -0.11 | 0.82 | 0.93 | 0.22 |
| *isw1* | 1 | 159.6 | 0.37 | -0.07 | 0.86 | 0.70 | 0.53 |
| *isw1* | 2 | 159.9 | 0.35 | -0.08 | 0.85 | 0.76 | 0.49 |
| *isw1* | 3 | 162.4 | 0.29 | -0.06 | 0.83 | 0.80 | 0.35 |
| *isw1* | 4 | 165.2 | 0.27 | -0.06 | 0.76 | 0.83 | 0.24 |
| *isw1* | 5 | 168.8 | 0.21 | -0.03 | 0.71 | 0.93 | 0.14 |
| *chd1* | 1 | 159.8 | 0.36 | +0.01 | 0.80 | 0.82 | 0.39 |
| *chd1* | 2 | 160.7 | 0.31 | -0.01 | 0.80 | 0.87 | 0.35 |
| *chd1* | 3 | 162.8 | 0.27 | -0.01 | 0.77 | 0.91 | 0.23 |
| *chd1* | 4 | 168.6 | 0.20 | -0.01 | 0.79 | 0.95 | 0.15 |
| *chd1* | 5 | 172.5 | 0.16 | 0.00 | 0.68 | 1.02 | 0.06 |
| *isw1 chd1* | 1 | - | 0.23 | +0.13 | 0.46 | 1.04 | 0.12 |
| *isw1 chd1* | 2 | - | 0.20 | +0.08 | 0.31 | 1.06 | 0.05 |
| *isw1 chd1* | 3 | - | 0.12 | +0.08 | 0.49 | 1.08 | 0.02 |
| *isw1 chd1* | 4 | - | 0.11 | +0.07 | 0.41 | 1.10 | 0.01 |
| *isw1 chd1* | 5 | - | 0.05 | +0.01 | 0.90 | 1.13 | 0.00 |

Depletion of Isw1 results in decreasing phasing amplitude and increasing spacing with increasing mean methylation rate (Q1 to Q5). These trends parallel those observed in wild type cells (Fig. 5B). Depletion of Chd1 shows similar trends to wild type and Isw1-depleted cells (Fig. 5C). In cells depleted of both Isw1 and Chd1, the mean methylation rate (Q1 to Q5) is always higher than in wild type and the single-degron mutants. Most strikingly, for cells depleted of both Isw1 and Chd1, the sine wave fit is very poor, with only a hint of nucleosome phasing, which almost disappears in Q5 (Fig. 5D). In particular, the +1 nucleosome is virtually undetectable in Q5, indicating that the positioning of the first nucleosome on these genes is essentially random. Thus, the fastest methylating genes in vivo have very little chromatin organization in the absence of Isw1 and Chd1.

### TFIIIB and TFIIIC transcription factor dynamics in vivo

Unlike nucleosomes, the binding of sequence-specific transcription factors to DNA is generally reversible on a seconds to minutes time scale in vitro and in vivo (reviewed by (*37*)). This means that detection of transcription factor binding is difficult in vitro unless binding site occupancy is very high (e.g., if the transcription factor concentration is very high, as in footprinting experiments), or if the salt concentration is very low (e.g., in the gel in a gel-shift assay) (see (*38*) for more discussion of this point). In vivo, rapidly reversible transcription factor binding has been demonstrated by FRAP experiments and single-molecule tracking (*37, 39, 40*). Consequently, it is unlikely that we can unambiguously detect the binding of typical transcription factors in our M.SssI methylation data. However, the binding of transcription factors TFIIIB and TFIIIC to tRNA genes and to other genes transcribed by RNA polymerase III (Pol III), is very stable in vitro and in nuclei (*20, 41, 42*). TFIIIB is composed of three subunits (including TATA-binding protein, TBP) and binds to the tRNA gene promoter, whereas TFIIIC is composed of six subunits and binds to two specific sites within the tRNA gene, termed the A and B boxes (Fig. 6A) (reviewed by (*43*)). The yeast genome contains 275 tRNA genes.

**Fig. 6.**
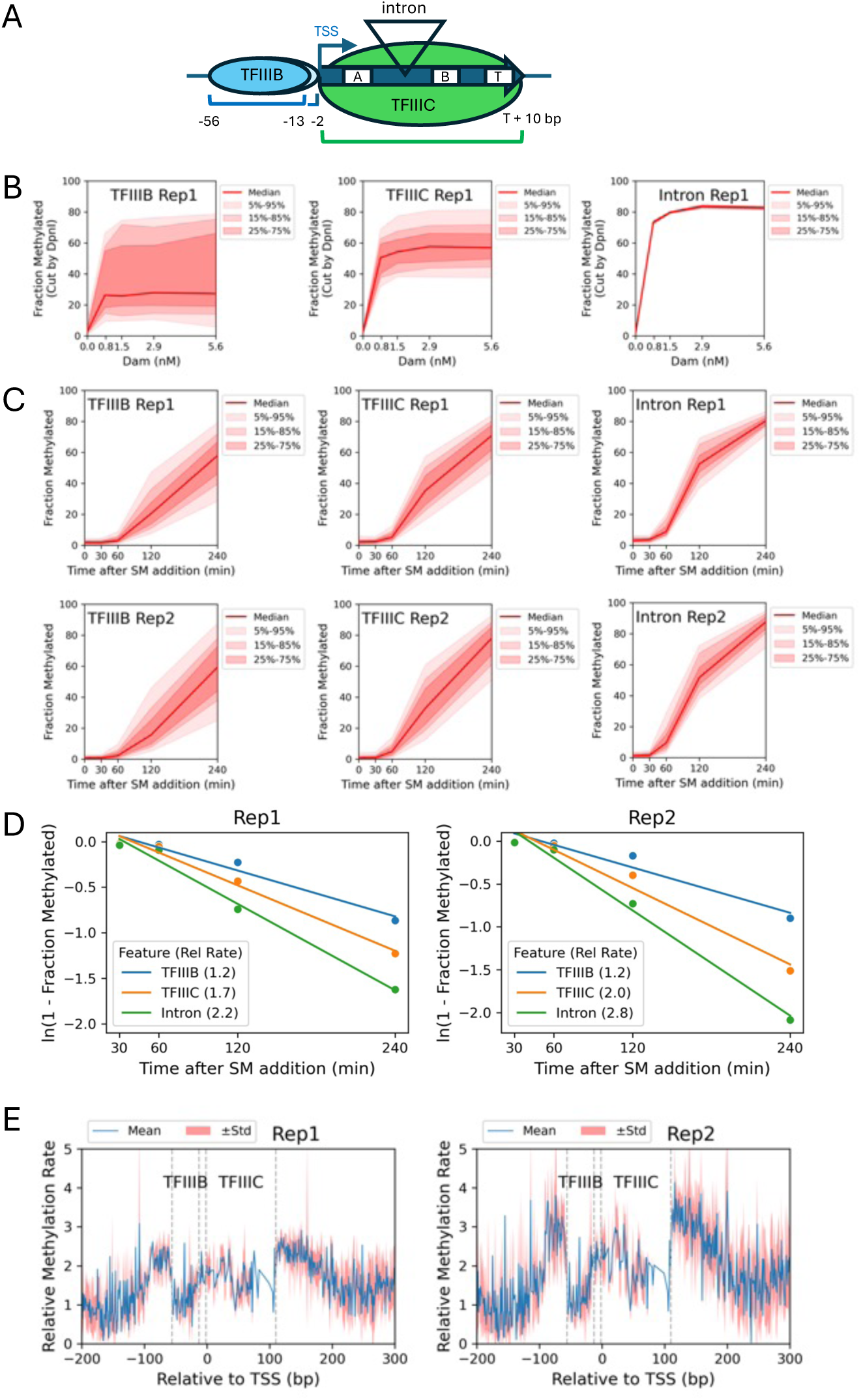
Dynamics of the TFIIIB and TFIIIC transcription factors at tRNA genes in nuclei and in vivo. (**A**) TFIIIB and TFIIIC binding at tRNA genes. (**B**) Dam methylation of GATC sites located in TFIIIB binding sites, TFIIIC binding sites and tRNA gene introns in nuclei, as a function of Dam concentration. Red line and shading: median GATC site methylation with data range shown. (**C**) Time courses after SM induction of M.SssI methylation of CG sites located in TFIIIB binding sites, TFIIIC binding sites and tRNA gene introns in vivo. (**D**) Methylation rates (normalized to the genomic median rate) for the median CG site located in TFIIIB binding sites, TFIIIC binding sites and tRNA gene introns in vivo (using data in C). (**E**) Methylation rate maps for the 215 tRNA genes without introns, aligned on the tRNA TSS. The TFIIIB and TFIIIC binding sites are indicated by dashed lines. The average methylation rate is depicted by the blue line; the standard deviation is depicted by the shaded area. Most tRNA genes are about 72 bp, but some are longer; consequently, the alignment results in a discontinuity in the data in the +72 to +99 region of the plot. The downstream region begins at +100.

Previously, we footprinted the TFIIIB-TFIIIC-tRNA transcription complex in isolated yeast nuclei using MNase-seq (*42*). We detected a stable complex with borders at -56 relative to the tRNA gene start site and at 10 bp downstream of the tRNA end (the Pol III terminator is also protected). We also observed two MNase cut sites between TFIIIB and TFIIIC, located at -13 and -2 relative to the tRNA gene start site. Here we define the TFIIIB binding site as -56 to -13, and the TFIIIC binding site as -2 to the end of the tRNA gene plus 10 bp, excluding the intron if present. Some tRNA genes have a single intron, which is located between the A and B boxes and loops out from the complex (Fig. 6A), where it is only weakly protected by TFIIIC from MNase digestion (*42*). We have also shown that tRNA genes have limited accessibility to Dam in nuclei, but are rapidly methylated in vivo (*20*).

We dissected the transcription complex into three regions: TFIIIB-bound, TFIIIC-bound and introns (Fig. 6A). We note that not all of the 275 tRNA genes in the yeast genome have GATC sites. In Dam-treated nuclei, the median limit methylation for the 22 GATC sites in TFIIIB-bound regions is less than that of the 133 GATC sites in TFIIIC-bound regions (∼27% and ∼57% methylated, respectively; Fig. 6B). This observation suggests that TFIIIB has higher average occupancy at tRNA genes than TFIIIC, in nuclei. There are only two intronic GATC sites, both of which are in the same gene (tP(UGG)O3, which specifies a prolyl tRNA). These two intronic sites are accessible to Dam in most nuclei (only protected in ∼20% of nuclei), but may not be representative of all tRNA gene introns. In conclusion, both TFIIIB and TFIIIC are stably bound in a fraction of nuclei, since methylation reaches a limit, but TFIIIB has a higher occupancy than TFIIIC.

We used the higher resolution afforded by the M.SssI data to assess the dynamics of TFIIIB and TFIIIC binding to tRNA genes in vivo. In total, tRNA genes contain 255 TFIIIB-bound CG sites, 1647 TFIIIC-bound CG sites and 54 intronic CG sites. All three sets of sites are rapidly methylated in vivo (Fig. 6C). Pseudo-first order rate plots give the expected straight lines with *R^2^* ≥ 0.94 (Fig. 6D). Methylation of TFIIIC binding sites is ∼1.5 times faster than methylation of TFIIIB binding sites, whereas intronic sites are methylated about twice as fast as TFIIIB sites (Fig. 6D). These data suggest that TFIIIB and TFIIIC occupancies in vivo mirror those in nuclei, with the critical difference that these factors are not stably bound in vivo, but dynamic, with different residence times, allowing internal GATC or CG site methylation.

We aligned the 215 tRNA genes without introns on their start sites and plotted the relative methylation rate at each CG site across tRNA genes (Fig. 6E). The methylation rate is lowest over most of the TFIIIB-bound promoter (from -56 to -13) and higher over the tRNA gene itself, where TFIIIC binds, consistent with TFIIIB being less dynamic than TFIIIC. The methylation rate is highest in the ∼50 bp just upstream and just downstream of tRNA genes. In conclusion, TFIIIB occupancy is higher than TFIIIC occupancy in vivo, as it is in nuclei. Although tRNA gene introns are relatively exposed, as in vitro, TFIIIB and TFIIIC do not protect tRNA genes strongly in vivo, perhaps due to displacement during Pol III transcription, which is not occurring in isolated nuclei.

## Discussion

A comparative analysis of methylation kinetics in remodeler mutants is now possible because Dam also methylates mtDNA in vivo. We speculate that Dam contains a serendipitous mitochondrial localization signal, although we were unable to identify one. Some potential confounding factors include indirect effects of remodeler loss on transport of Dam into mitochondria or on SAM concentration. However, the use of mtDNA to normalize the data for differences in Dam concentration and induction kinetics is supported by the similar mtDNA methylation rates observed in the biological replicate experiments (Fig. 1D), and by the close agreement between the phasing profiles for the biological replicates after normalization (Fig. 2).

Rsc8-depleted cells methylate their DNA at about the same rate as wild type cells, suggesting that RSC plays a minimal role in driving nucleosome dynamics. During the induction time course, the nucleosomes are shifting to new average positions closer to the promoter (*20*), consistent with MNase-seq data for nuclei from cells completely depleted of Rsc8 (*25, 26*). Although the nucleosome shifts show that Rsc8 depletion is effective, there are some important caveats. These are: (i) Rsc8 is not completely depleted until the end of the time course; (ii) the cells are arresting because Rsc8 is essential; (iii) loss of Rsc8 may release the Sth1-Arp7-Arp9-Rtt102 ATPase module from the complex, which can remodel nucleosomes in vitro (*44*), such that RSC-dependent nucleosome dynamics may continue rather than cease, presumably in an unregulated mode.

Isw1-depleted cells and Chd1-depleted cells also methylate their DNA at about the same rate as wild type cells. Chd1 is depleted very rapidly, but Isw1 depletion is slower; neither is essential for growth. However, cells depleted of both Isw1 and Chd1 methylate their DNA faster than wild type cells. The major disruption in nucleosome organization expected from MNase-seq data for the double null mutant (*17, 18*) is also apparent, indicating that extensive nucleosome mobilization occurs as the remodelers are depleted in vivo (*20*). The increased methylation rate could be explained by increased nucleosome mobilization, or by some nucleosome loss resulting in longer linkers of variable length, or by increased levels of conformationally altered nucleosomes (*20*). Support for the nucleosome loss model is provided by evidence that Isw1 and Chd1 prevent histone exchange (*45*). As a working model, we propose that the ISW1 and CHD1 remodeling complexes together suppress nucleosome dynamics in vivo, by reversing the effects of disruptive remodeling activities to re-create an ordered static array, as in isolated nuclei, and that yeast genes cycle between remodeled and ordered states (Fig. 7).

**Fig. 7.**
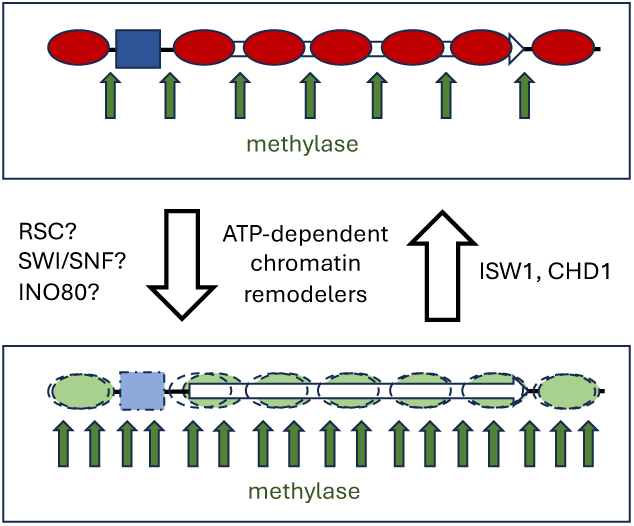
Working model for dynamic chromatin in vivo. Remodeled nucleosomes may be displaced, slid along the DNA, and/or conformationally altered. Genes cycle between an ordered static state as in nuclei (top) and a remodeled dynamic state (bottom), due to opposing remodeling activities. ISW1 and CHD1 suppress nucleosome dynamics by converting remodeled chromatin to static canonical chromatin. RSC, SWI/SNF and/or INO80 may be responsible for creating dynamic remodeled chromatin. The promoter is shown occupied by a transcription complex which cycles on and off the DNA in vivo (blue box).

We have presented a decaying sine wave model to quantify nucleosome phasing profiles using five parameters. The amplitude is a measure of the degree of phasing, such that larger amplitude indicates better phasing, as more nucleosomes are positioned closer to the average position. The wavelength indicates the average nucleosome spacing. The decay factor measures the degree to which the phasing signal fades as a function of distance from the promoter. The change in mean methylation rate over 1 kb is measured by the slope of the mean methylation rate as a function of distance from the promoter.

Finally, the mean fit is the average methylated fraction or methylation rate for the first kilobase from the +1 nucleosome dyad. The fit of the sine wave model is excellent for wild type cells, although it is weaker for the remodeler mutants as their nucleosomal arrays are less ordered. Nevertheless, the model provides reliable measures of the important parameters describing phased nucleosome arrays.

For nuclei, phasing profiles can be fit to the limit methylation state (i.e., when the fraction methylated does not change with increasing enzyme concentration). However, for living cells, the amplitude changes with time as the DNA becomes fully methylated (*20*). We resolved this issue by calculating the methylation rate at each GATC or CG site and plotting this parameter against distance from the +1 nucleosome dyad. The methylation rate was used to quantify the relative accessibility of the entire genome.

Intriguingly, the methylation rate decreases with distance from the promoter in wild type cells in vivo. We interpret this to mean that nucleosome dynamics decrease with distance from the promoter. That is, the +1 nucleosome is more dynamic than the +2 nucleosome, etc. However, the fact that this effect is observed in isolated nuclei, albeit much weaker, suggests that there is a stable component to the methylation rate decrease along genes. Potential explanations for decreasing methylation rate with distance from the promoter include faster histone turnover in promoter-proximal nucleosomes (*45–47*), higher levels of H2A.Z in promoter-proximal nucleosomes (*48*), and changes in various histone marks along genes, such as methylation and acetylation (*49*). Histone turnover would only affect the methylation rate in vivo because it is an ATP-dependent process, but H2A.Z and histone marks may also affect the methylation rate in nuclei, since they are preserved after isolation. For example, acetylated histone tail domains bind less tightly to DNA and increase the association/dissociation rates of terminal nucleosomal DNA (*50*), potentially increasing the methylation rate in nuclei.

The decreasing methylation rate with distance from the promoter is less pronounced in the absence of Isw1 and almost disappears in the absence of Chd1. The interaction of Chd1 with H3K4me3 (*51*) suggests a direct explanation for the requirement for Chd1, since H3K4 methylation also decreases with distance from the promoter (*49, 52, 53*). Similarly, Isw1 binding to some active genes is mediated by H3K4me3 (*54*), although the nucleosome spacing activity of Isw1 is not globally dependent on H3K4me3 (*55*). In cells depleted of both Isw1 and Chd1, the Dam methylation rate actually increases with distance from the promoter. An additional factor in this double mutant is extensive di-nucleosome enrichment involving the +1, +2 and +3 nucleosomes (*56*); the absence of a linker would result in slower methylation nearer the promoter, although the overall methylation rate is faster than in wild type and the single mutants in vivo.

We have also gained some insight into transcription factor dynamics in vivo. TFIIIB and TFIIIC are not typical transcription factors because they are large multi-subunit complexes that are stably bound to tRNA genes in vitro. However, they are highly dynamic in vivo, possibly resulting from displacement of both transcription factors each time Pol III transcribes a tRNA gene, transiently exposing the DNA to the methylase.

## Materials and Methods

### Experimental Design

Our aim is to assess the roles of RSC, ISW1 and CHD1 in nucleosome dynamics in living cells. Yeast strains were constructed with an auxin-dependent degron attached to the gene encoding a subunit of one of these chromatin remodeling complexes. These strains also contained an SM-inducible gene for expression of either the Dam or the M.SssI DNA methyltransferase. Our initial analysis of data for Rsc8 depletion and depletion of both Isw1 and Chd1 was described previously (*20*). Here, we add the Isw1-degron and Chd1-degron single mutants and discover that Dam also methylates mtDNA in vivo. Since mtDNA is non-nucleosomal, we make the assumption that the mtDNA methylation rate is determined by the Dam concentration and is independent of chromatin structure. This approach allows us to compare methylation rates between wild type and remodeler degron mutants in living cells.

### Yeast Strains

Isw1 degron (YHP820) and Chd1 degron (YHP821) strains were obtained by transformation of strain YHP808 (*MATa ade2-1 can1-100 leu2-3,112 trp1-1 ura3-1*, *hoΔ::TIR1-tetO2 tCUP(1xGcn4)-Dam-Deg-3HA KanMX*) with NotI digests of plasmids p892 and p893, respectively. Detailed descriptions of the construction of these plasmids and the wild type (YHP808), Rsc8-degron (YHP812) and Isw1/Chd1 double degron (YHP823) strains, can be found in (*20*).

### Immunoblotting

Dam-degron-3HA, Isw1-degron-3HA, and Chd1-degron-3HA were detected using horse radish peroxidase-conjugated anti-HA antibody (3F10; Roche 12013819001). Tubulin was used as a loading control and detected with horseradish peroxidase-conjugated anti-tubulin antibody (Abcam ab-185067). Immunoblotting was performed as described (*20*).

### Computational Analysis

We processed Dam methylated data using a snakemakeMethylFrac pipeline (https://github.com/zhuweix/snakemakeMethylFrac) (*22*). This pipeline does the following: 1) The raw Illumina sequencing data (GSE230309) (*20*) are aligned to the sacCer3 genome using bowtie2 v2.5.1 (*57*). 2) The read fragment occupancy and 5’-ends are counted using bedtools v2.31.1 (*58*). GATC half sites closer than 200 bp were filtered out because DNA fragments < 200 bp tend to be lost during sample preparation. 3) The methylated fractions for each GATC site are stored in bigwig files using pyBigWig 0.3.22 (https://github.com/deeptools/pyBigWig).

Bases in the raw M.SssI nanopore sequencing data were called using dorado v0.5.0 (https://github.com/nanoporetech/dorado) with DNA model sup v3.3, 5mCG_5hmCG. The reads were aligned to sacCer3 using dorado. The methylated fraction of each CG site (5hmCG was excluded) was calculated using modkit v0.4.1 (https://github.com/nanoporetech/modkit) with parameters --cpg --combine-strands -- ignore h. Methylated fractions are stored in bigwig files created using pyBigWig 0.3.22.

We performed the RNAseq analysis using two biological replicates of yeast treated with SM for 25 min in GSE110413 (*36*) using salmon 1.10.1 (*59*) and calculated the average transcription level in Total Per Million (TPM). MNase phasing data were obtained from (*18*).

We developed python software to perform the downstream analysis (https://github.com/zhuweix/SinePhasingAnalyzer). The software combines the data in the bigwig files into an SQLite database and calculates the percentiles of features, the average phasing relative to the +1 nucleosome dyads of protein-coding genes (smoothed using 21-bp windows), the phasing relative to the TSS for tRNA genes, the methylation rates for each GATC or CG site, and performs the decaying sine wave analysis. This software used NumPy (*60*), pandas (*61*), pyBigWig, matplotlib (*62*), seaborn (*63*), SciPy (*64*) and statsmodels (*65*).

We developed a decaying sine wave model (formula 1). The amplitude is assumed to decay exponentially and to reach 0 downstream of the gene body. ‘*x*’ is the location relative to the dyad of the +1 nucleosome. The nucleosome spacing is the period of a sine wave,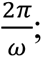 the amplitude at the +1 nucleosome dyad is *A*; the slope, *k*, is the change in the baseline of the sine wave and 1000*k* is the change in baseline over 1 kb; the baseline at the +1 nucleosome dyad is *b*; the proportional amplitude decrease after one period (decay) is 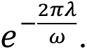 The adjusted average is the average of the fitted curve. The single-gene analysis takes into account that the CG sites in a gene are unevenly distributed; the simple mean of the methylation rates is biased due to the CpG distribution. We used the fitted curve for all yeast genes to estimate the bias. We calculated the bias of each gene as the actual average methylation rate of the CG sites relative to the values from the fitted curve for all genes. Thus, we computed the unbiased average relative methylation rate as average methylation rate – methylation rate bias estimated from the fitted curve.

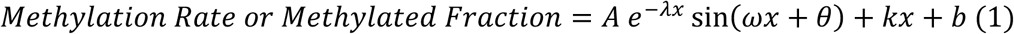

### Statistical Analysis

The methylation rate of each GATC or CG site was estimated using first-order kinetics in formula 2. The methylated fractions were capped at 0.99. If multiple data points were > 0.99, only the first of these points was preserved to avoid saturation. The apparent rate *k* was computed using linear regression and *R^2^* was calculated. The methylation rate of mtDNA was calculated using the median methylated fractions, and used to normalize the methylation rates amongst strains for Dam methylated samples. The M.SssI methylated samples were normalized to the median genome methylation rate calculated from the median methylated fraction for all 16 yeast chromosomes, mtDNA and 2-µm plasmid.

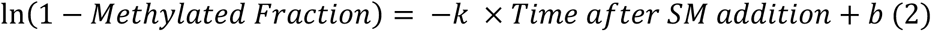

The adjusted *R^2^*of the sine wave fit was computed using formula 3, where SSR is the sum of squared residuals, SST is the total sum of squares, and N is the number of GATC or CG sites for curve fit (N=1051 for locations from -50 to 1000 relative to the +1 nucleosome dyad).

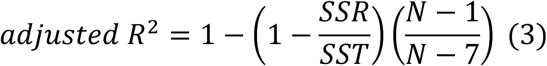

We developed an interactive web application, Phasing Analysis App, which is hosted on https://sinewaveanalyzer.streamlit.app/. This app performs decaying sine wave phasing analysis using the raw phasing data and allows the user to test the influence of each parameter on the fitted curve.

## Acknowledgments

This study utilized the high-performance computational capabilities of the Biowulf Linux cluster at the National Institutes of Health (NIH).

## Author contributions

Conceptualization: ZX, DJC

Methodology: ZX, HKP

Investigation: ZX, HKP, PRE

Visualization: ZX, HKP, DJC

Supervision: DJC

Writing—original draft: DJC

Writing—review & editing: DJC, ZX, HKP, PRE

## Data and materials availability

The Illumina and Nanopore sequence data described in this paper are publicly available at the GEO database under accession number GSE230309. The sacCer3 genome sequence is available at the Saccharomyces Genome Database. Bigwig files containing methylated fraction data are available at figshare.

All data are available in the main text or the supplementary materials.

The snakemakeMethylFrac pipeline is available at https://github.com/zhuweix/snakemakeMethylFrac

The Sine Phasing Analyzer is available at https://github.com/zhuweix/SinePhasingAnalyzer

The web application Phasing Analysis App is available at https://sinewaveanalyzer.streamlit.app/

## Supplementary Materials

**Fig. S1.**
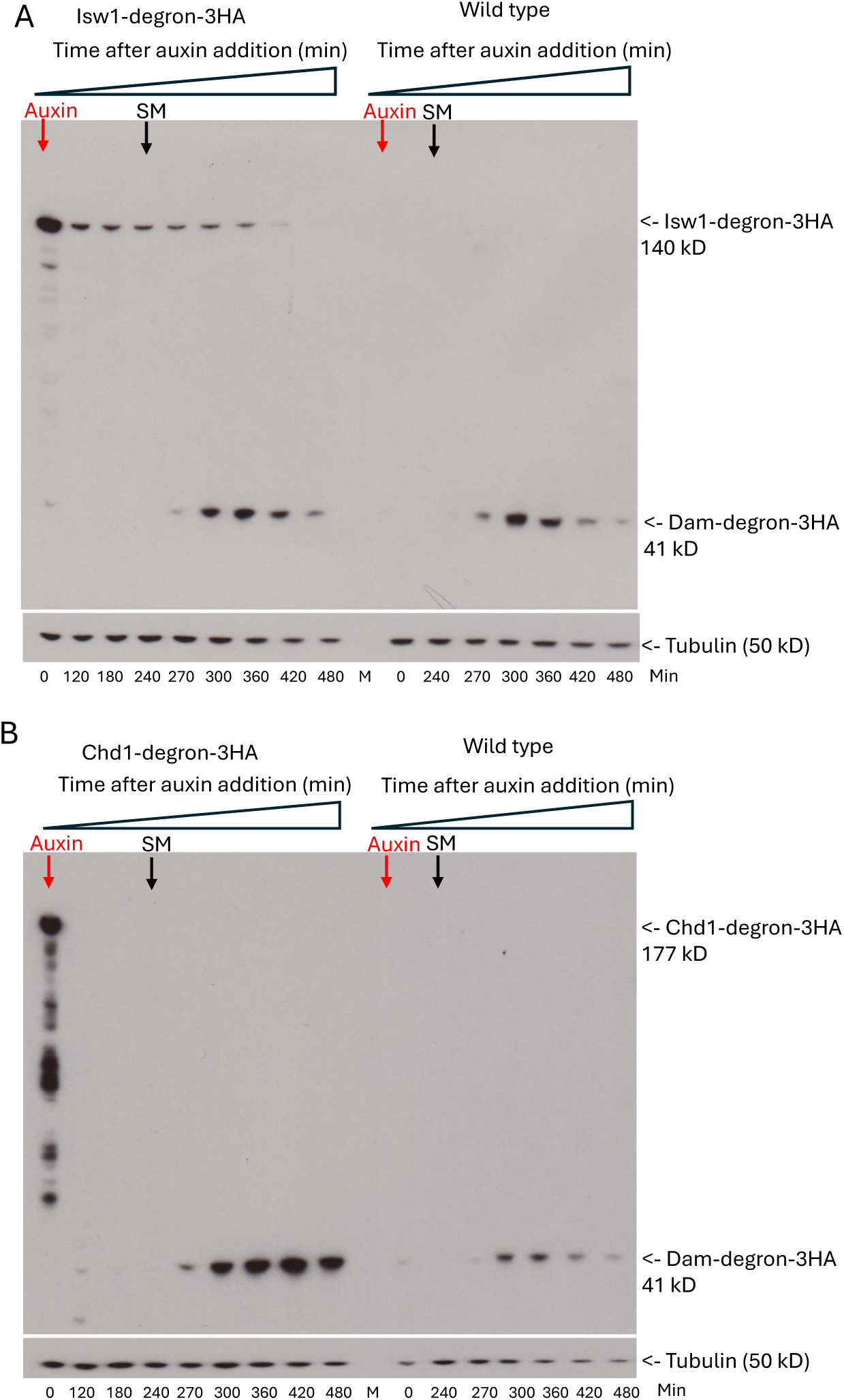
Chd1-depleted cells produce more Dam than wild type cells. **(A)** Anti-HA immunoblot performed to follow Isw1 depletion and Dam induction. Wild type and Isw1–degron cells were treated with auxin for 4 h and then induced with SM for another 4 h. Our analysis begins with the 240 min time point, when SM was added (= 0 min). (**B**) Anti-HA immunoblot showing higher Dam induction compared to wild type after depletion of Chd1.

**Fig. S2.**
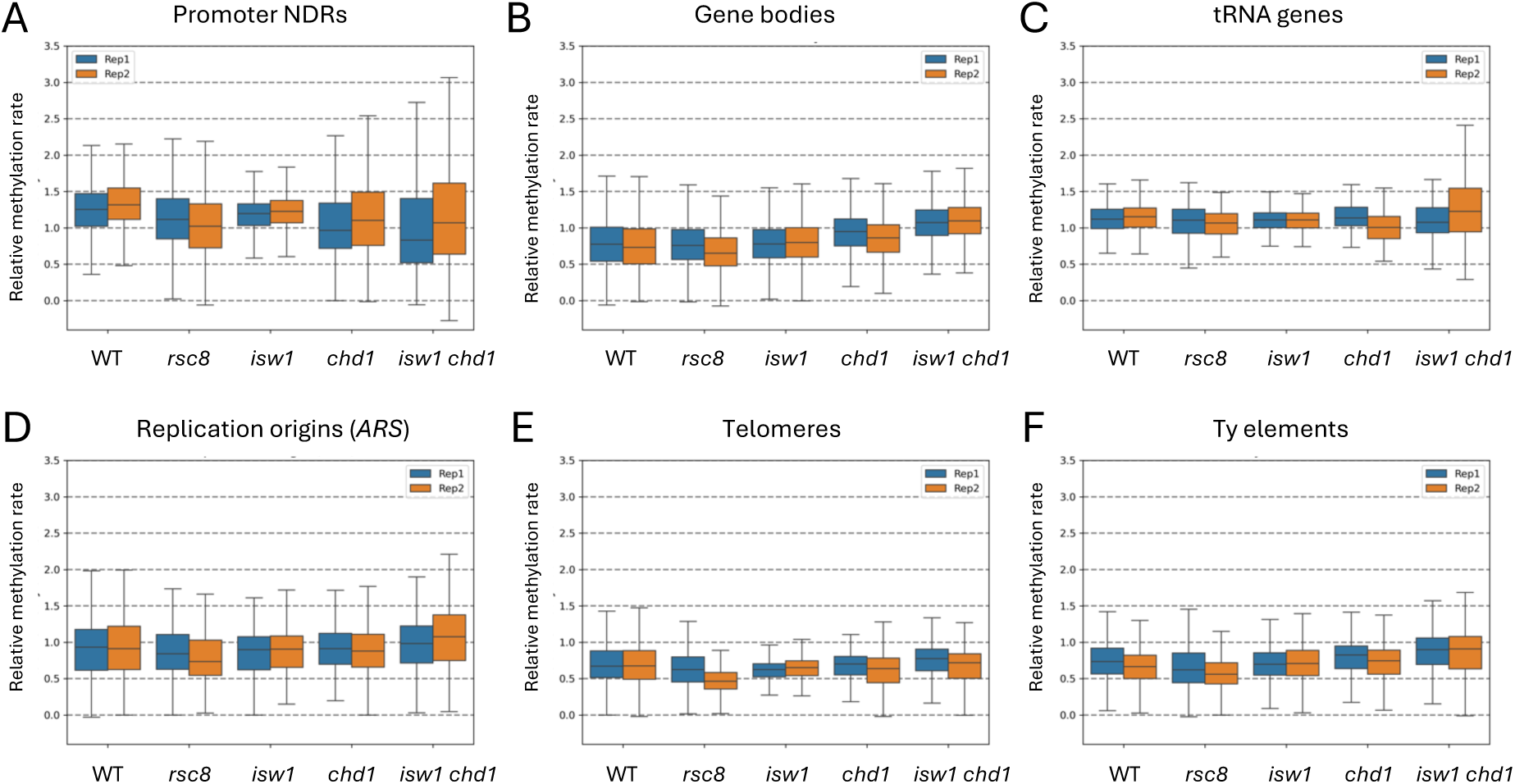
Comparison of normalized average Dam methylation rates in vivo in various genomic elements: wild type vs. cells depleted of remodeler subunits. The methylation rate was calculated for each GATC site in the yeast genome and normalized to the internal median mtDNA methylation rate (set to 1). (**A**) Promoter NDRs. (**B**) Gene bodies. (**C**) tRNA genes. (**D**) Replication origins (*ARS* elements). (**E**) Telomeres. (**F**) Ty transposable elements. The box contains 25%-75% of the data; the line is the median; the whiskers represent 1.5 times the inter-quartile range to the farthest data points.

**Fig. S3.**
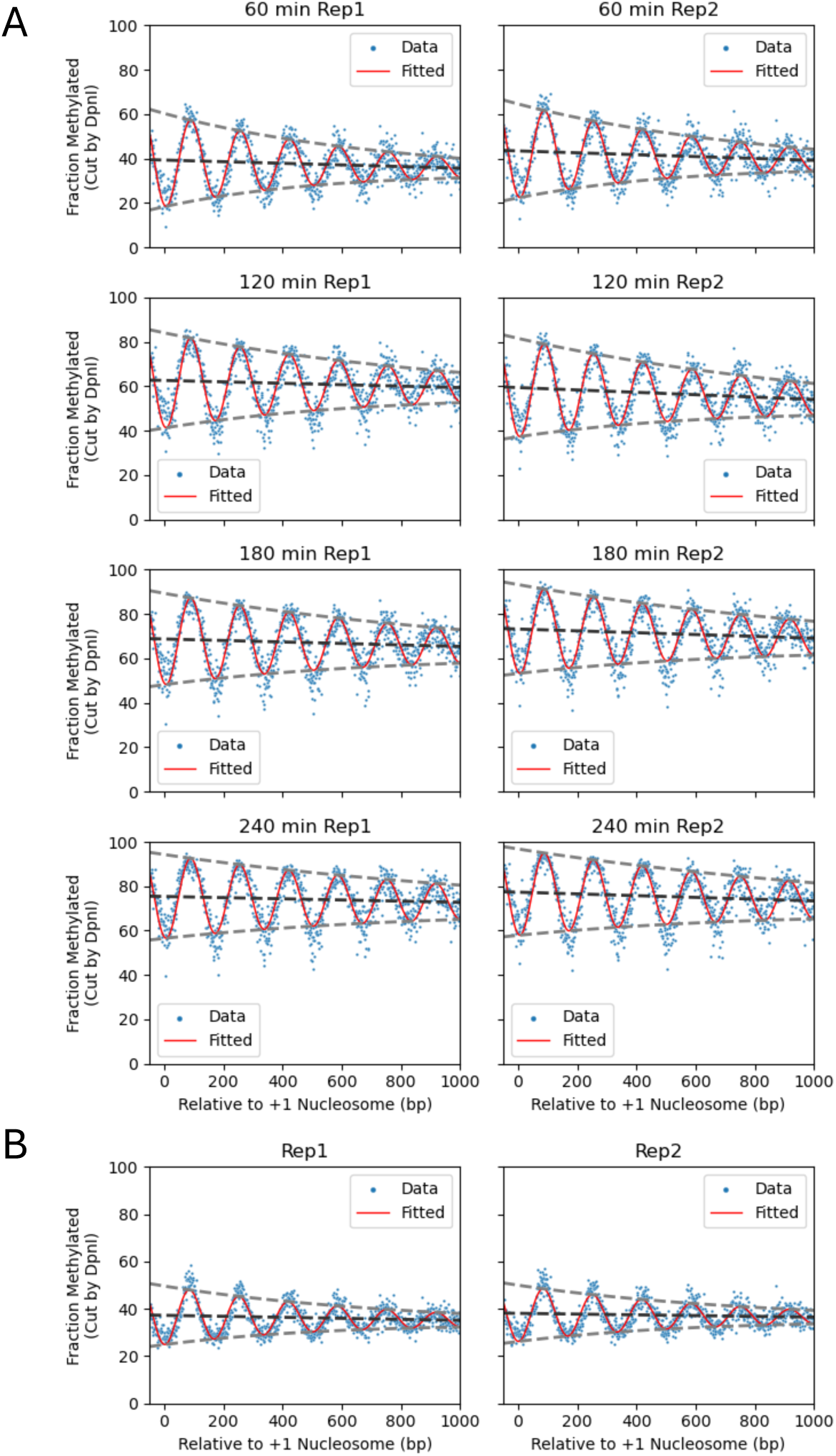
Decaying sine wave model fit to Dam methylation data for wild type, remodeler-depleted cells and for isolated nuclei. (**A**) Sine wave model fit to time course data for wild type cells. The methylated fraction was calculated for each GATC site in the yeast genome at each time point after SM addition. The methylated fraction is plotted relative to the location of the +1 nucleosome dyad of 5398 genes. Values for individual GATC sites are indicated by blue dots. The decaying sine wave fit is shown as a red line. The baseline of the sine wave is indicated by the dark grey dashed line; the decaying amplitudes are indicated by light grey dashed lines. (**B**) Analysis of limit methylation data for wild type nuclei treated with 5.6 nM Dam.

**Fig. S4.**
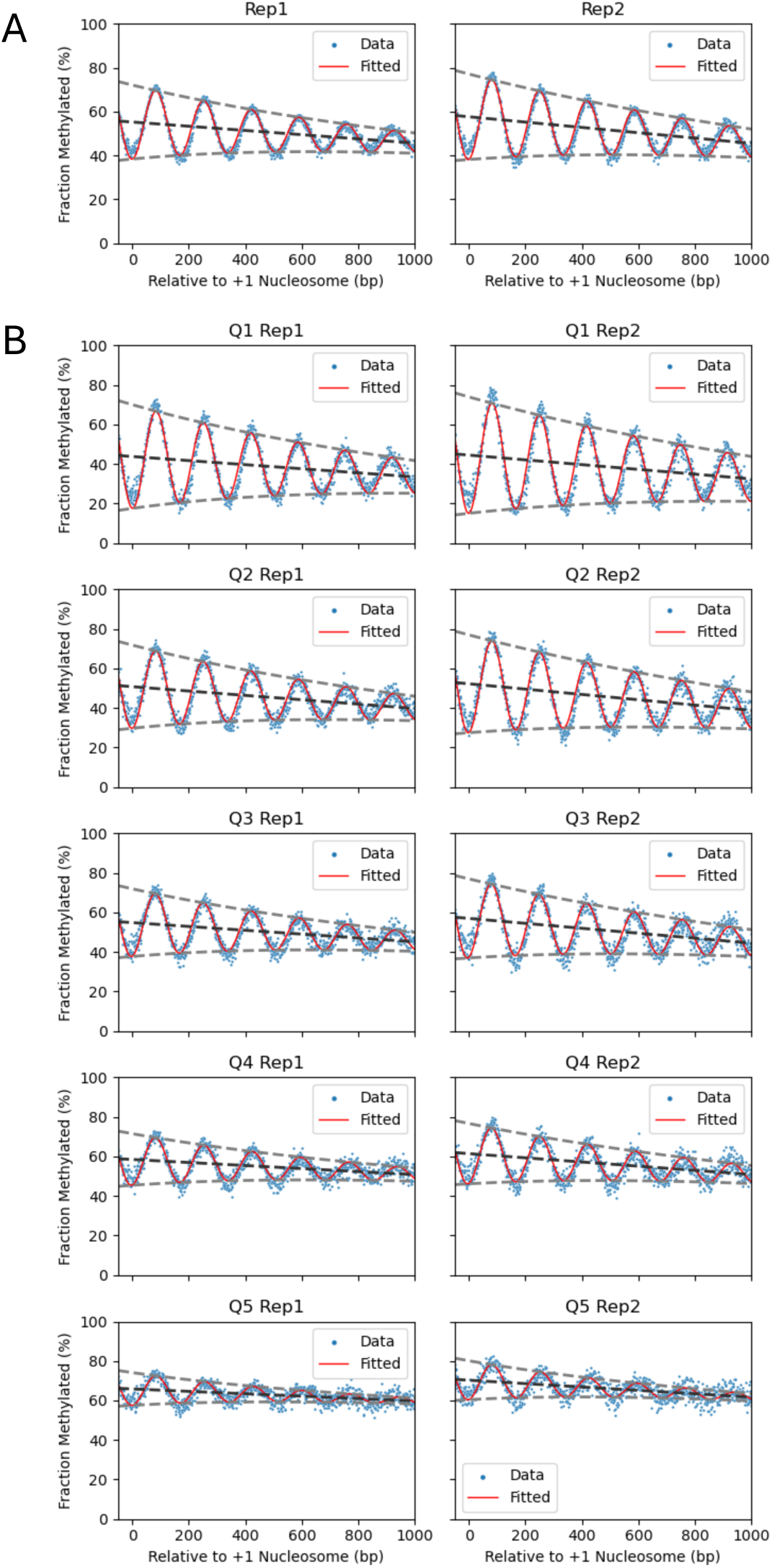
Decaying sine wave model fit to M.SssI methylation nanopore data for wild type cells induced with SM for 240 min in vivo. (**A**) Analysis of all genes. The methylated fraction was calculated for each CG site in the yeast genome. Individual CG sites are indicated by blue dots. The decaying sine wave fit is shown as a red line. The baseline of the sine wave is indicated by the dark grey dashed line; the decaying amplitudes are indicated by light grey dashed lines. **(B)** Analysis of genes sorted into quintiles using their calculated individual unbiased average methylation rates. Quintile 1 (Q1) contains the slowest methylating genes.

**Fig. S5.**
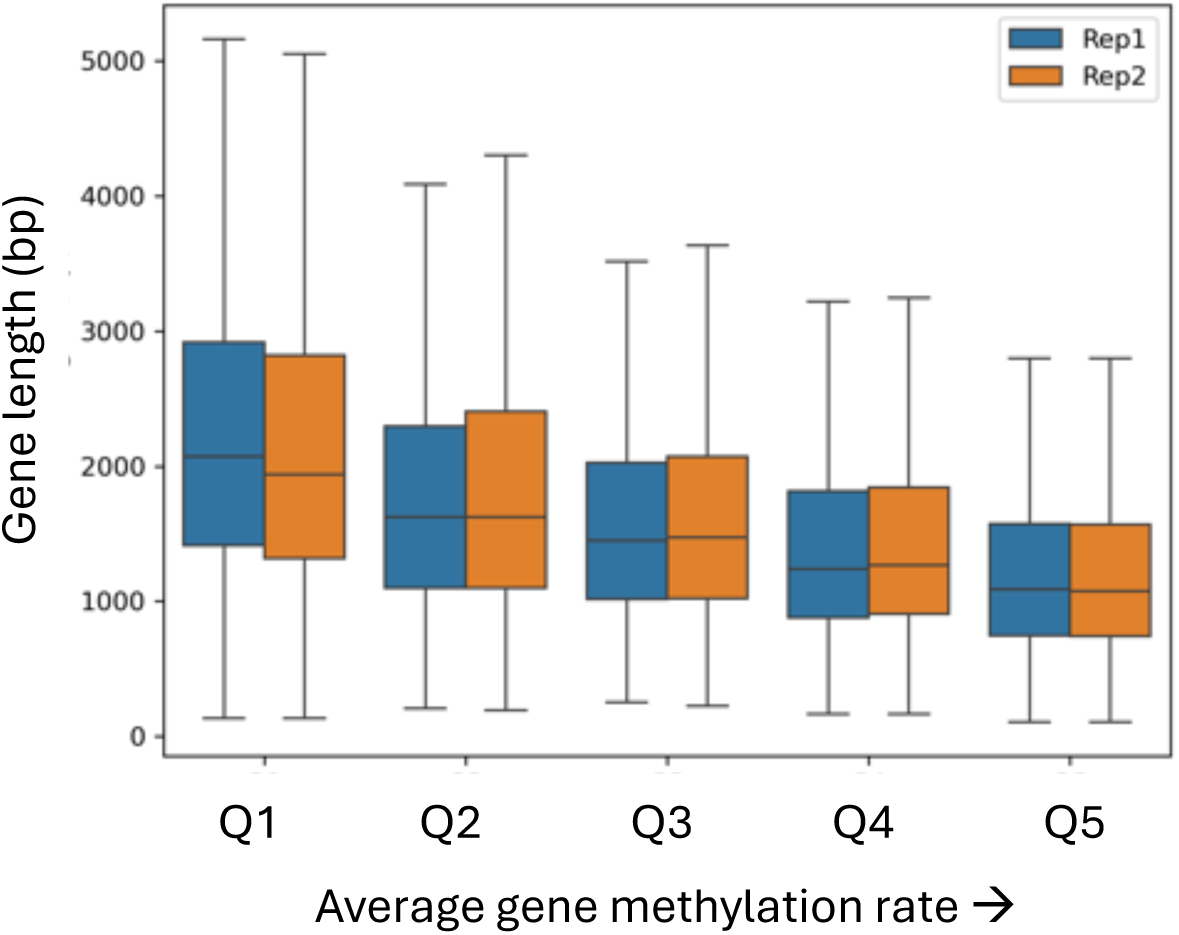
Gene length distributions in the mean relative M.SssI methylation rate quintiles. The 5398 genes were divided into quintiles according to their individual mean methylation rates, as shown in Fig. 4B. Quintile 5 (Q5) contains the fastest methylating genes. Box plots show the distribution of gene lengths (TSS to TTS) for the genes in each quintile. The box contains 25%-75% of the data; the line is the median; the whiskers represent 1.5 times the inter-quartile range to the farthest data points).

**Fig. S6.**
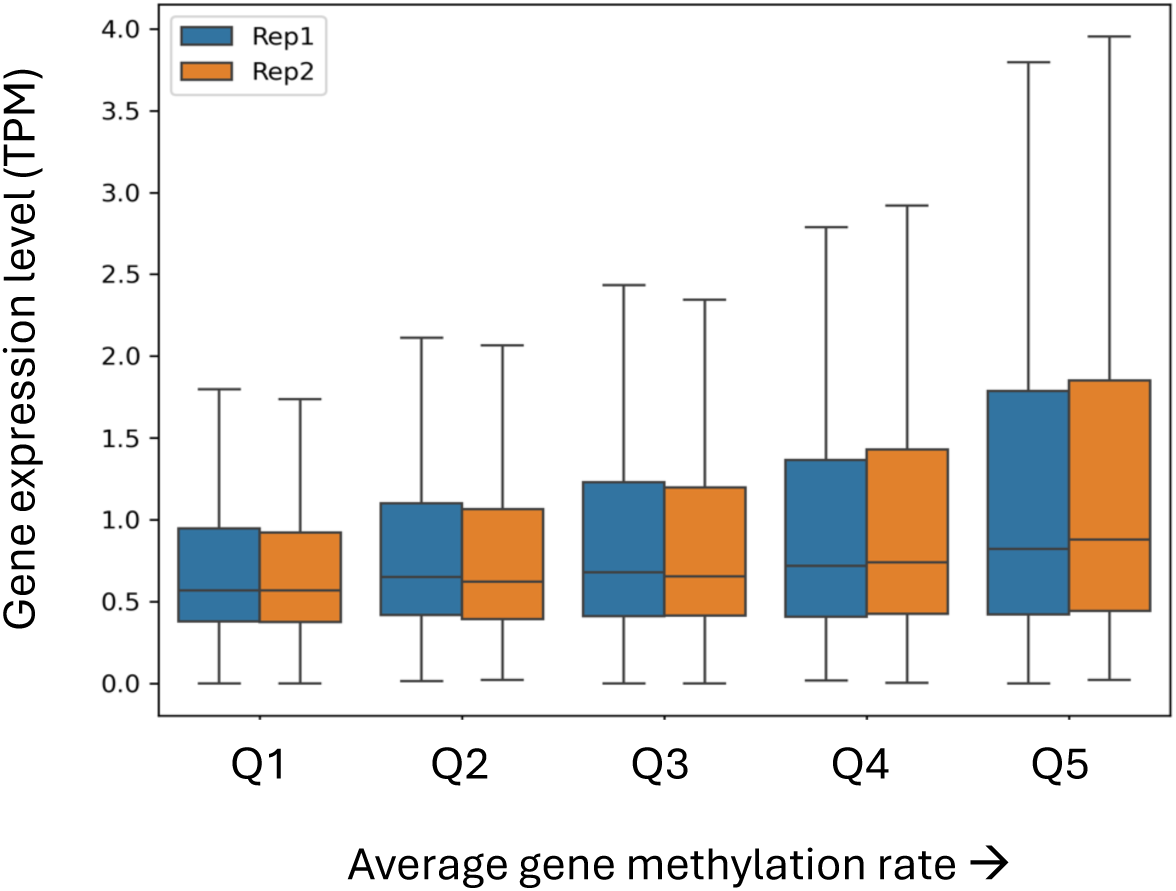
Transcript level distributions in the mean relative M.SssI methylation rate quintiles. The 5398 genes were divided into quintiles according to their individual mean methylation rates, as shown in Fig. 4B. Quintile 5 (Q5) contains the fastest methylating genes. Box plots show the distribution of transcript levels for the genes in each quintile (RNA-seq data for SM-induced cells (*33*)) for the genes in each quintile. The box contains 25%-75% of the data; the line is the median; the whiskers represent 1.5 times the inter-quartile range to the farthest data points).

**Fig. S7.**
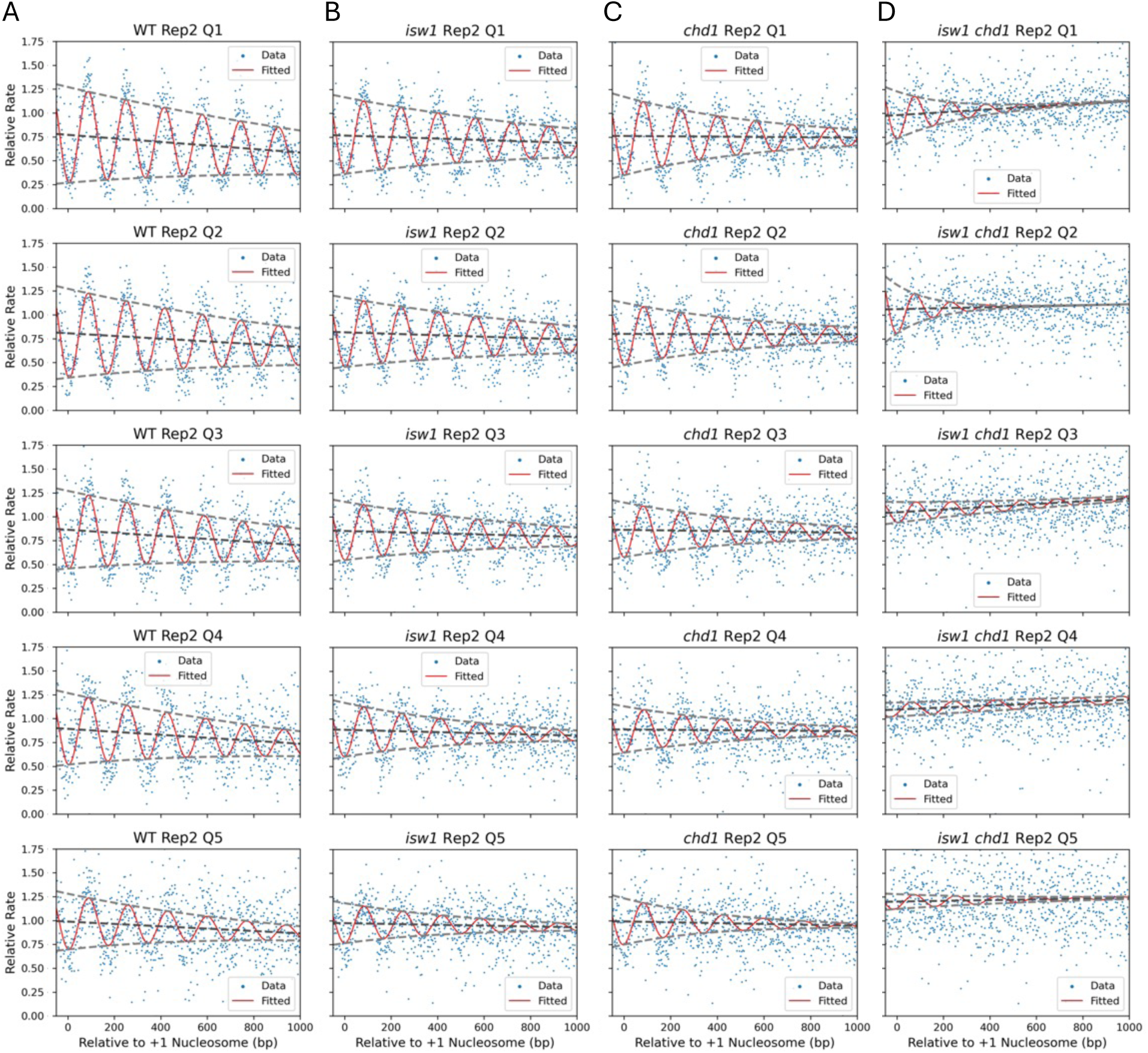
Decaying sine wave model fit to Dam methylation data for wild type and remodeler-depleted cells in vivo: Comparison of methylation rate quintiles for biological replicate 2. (**A**) wild type; (**B**) Isw1-degron mutant; (**C**) Chd1-degron mutant; (**D**) Isw1-degron and Chd1-degron mutant. (see Table S2 for fit parameters). Dam methylation rate data normalised to mtDNA (set to 1) are plotted for each quintile of genes defined using M.SssI nanopore data for replicate 1 (Fig. 4B). Values for individual GATC sites are indicated by blue dots. The methylation rates for each set of genes are plotted relative to the location of the +1 nucleosome dyad. The decaying sine wave fit is shown as a red line. The baseline of the sine wave is indicated by the dark grey dashed line; the decaying amplitudes are indicated by light grey dashed lines. Data for biological replicate 1 are shown in Fig. 5.

**Table S1.**
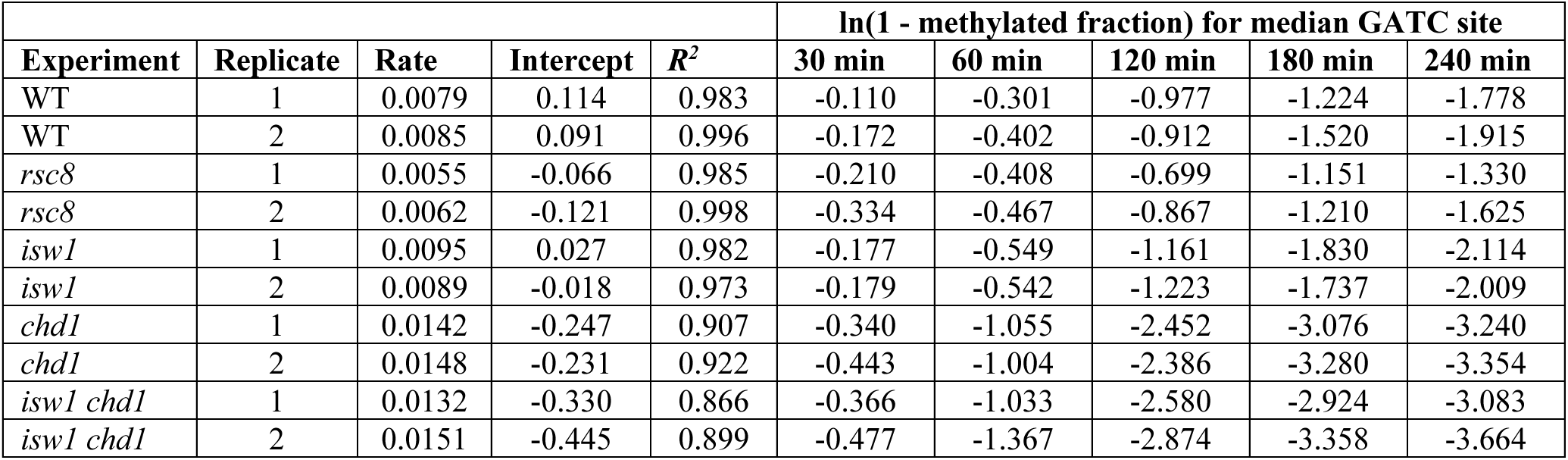
mtDNA methylation rates in vivo for wild type and remodeler depletion mutants. Calculation of ‘ln(1 - methylated fraction)’ at times after SM addition. The data for wild type (WT) mtDNA replicate 1 are plotted in Fig. 1C. The median methylation rate for the GATC sites in mtDNA is given in the ‘Rate’ column. The rate is given by the slope of the plot of ‘ln(1 - methylated fraction)’ vs. time after SM addition (the intercept and the coefficient of determination (*R^2^*) are also provided; formula 2). The distributions of the rate values for individual GATC sites in mtDNA for each replicate are shown in the box plots in Fig. 1D.

| Experiment | Replicate | Rate | Intercept | $R^2$ | ln(1 - methylated fraction) for median GATC site | | | | |
| --- | --- | --- | --- | --- | --- | --- | --- | --- | --- |
|  |  |  |  |  | 30 min | 60 min | 120 min | 180 min | 240 min |
| WT | 1 | 0.0079 | 0.114 | 0.983 | -0.110 | -0.301 | -0.977 | -1.224 | -1.778 |
| WT | 2 | 0.0085 | 0.091 | 0.996 | -0.172 | -0.402 | -0.912 | -1.520 | -1.915 |
| *rsc8* | 1 | 0.0055 | -0.066 | 0.985 | -0.210 | -0.408 | -0.699 | -1.151 | -1.330 |
| *rsc8* | 2 | 0.0062 | -0.121 | 0.998 | -0.334 | -0.467 | -0.867 | -1.210 | -1.625 |
| *isw1* | 1 | 0.0095 | 0.027 | 0.982 | -0.177 | -0.549 | -1.161 | -1.830 | -2.114 |
| *isw1* | 2 | 0.0089 | -0.018 | 0.973 | -0.179 | -0.542 | -1.223 | -1.737 | -2.009 |
| *chd1* | 1 | 0.0142 | -0.247 | 0.907 | -0.340 | -1.055 | -2.452 | -3.076 | -3.240 |
| *chd1* | 2 | 0.0148 | -0.231 | 0.922 | -0.443 | -1.004 | -2.386 | -3.280 | -3.354 |
| *isw1 chd1* | 1 | 0.0132 | -0.330 | 0.866 | -0.366 | -1.033 | -2.580 | -2.924 | -3.083 |
| *isw1 chd1* | 2 | 0.0151 | -0.445 | 0.899 | -0.477 | -1.367 | -2.874 | -3.358 | -3.664 |

**Table S2.** Decaying sine wave model parameters fitted to Dam data for wild type and depletion mutants in vivo. For the quintile data in Fig. S7 (Replicate 2): individual genes grouped by methylation rate (Q5 contains the fastest methylating genes). Parameters are as described in the legend to Table 1. Spacing values are not provided for the Isw1/Chd1 double depletion mutant because the fit is too poor. See Table 3 for the fit to biological replicate 1.

| Strain | Quintile | Spacing (bp) | Amplitude | Slope per kb | Decay per Period | Adjusted Mean Rate | Adjusted $R^2$ |
| --- | --- | --- | --- | --- | --- | --- | --- |
| WT | 1 | 163.9 | 0.50 | -0.18 | 0.88 | 0.67 | 0.66 |
| WT | 2 | 164.2 | 0.47 | -0.14 | 0.86 | 0.73 | 0.58 |
| WT | 3 | 166.4 | 0.41 | -0.16 | 0.87 | 0.78 | 0.47 |
| WT | 4 | 169.0 | 0.37 | -0.15 | 0.84 | 0.81 | 0.36 |
| WT | 5 | 170.3 | 0.29 | -0.12 | 0.80 | 0.93 | 0.22 |
| *isw1* | 1 | 159.9 | 0.40 | -0.08 | 0.85 | 0.72 | 0.54 |
| *isw1* | 2 | 160.6 | 0.36 | -0.08 | 0.86 | 0.77 | 0.50 |
| *isw1* | 3 | 162.7 | 0.31 | -0.06 | 0.83 | 0.82 | 0.36 |
| *isw1* | 4 | 164.9 | 0.28 | -0.06 | 0.77 | 0.85 | 0.25 |
| *isw1* | 5 | 167.8 | 0.21 | -0.04 | 0.75 | 0.95 | 0.14 |
| *chd1* | 1 | 160.0 | 0.41 | -0.02 | 0.78 | 0.75 | 0.42 |
| *chd1* | 2 | 160.4 | 0.33 | 0.00 | 0.79 | 0.79 | 0.32 |
| *chd1* | 3 | 164.3 | 0.29 | -0.03 | 0.78 | 0.84 | 0.24 |
| *chd1* | 4 | 168.2 | 0.24 | -0.01 | 0.76 | 0.88 | 0.16 |
| *chd1* | 5 | 171.8 | 0.24 | -0.05 | 0.67 | 0.97 | 0.09 |
| *isw1 chd1* | 1 | - | 0.25 | +0.15 | 0.55 | 1.05 | 0.12 |
| *isw1 chd1* | 2 | - | 0.26 | +0.05 | 0.39 | 1.08 | 0.07 |
| *isw1 chd1* | 3 | - | 0.11 | +0.15 | 0.80 | 1.12 | 0.05 |
| *isw1 chd1* | 4 | - | 0.07 | +0.11 | 0.88 | 1.15 | 0.02 |
| *isw1 chd1* | 5 | - | 0.08 | +0.04 | 0.72 | 1.22 | 0.00 |

